# Auditory dyadic interactions through the ‘eye’ of the social brain: How visual is the posterior STS interaction region?

**DOI:** 10.1101/2023.03.13.532398

**Authors:** Julia Landsiedel, Kami Koldewyn

## Abstract

Human interactions contain potent social cues that not only meet the eye but also the ear. Although research has identified a region in the posterior superior temporal sulcus as being particularly sensitive to visually presented social interactions (SI-pSTS), its response to auditory interactions has not been tested. Here, we used fMRI to explore brain response to auditory interactions, with a focus on temporal regions known to be important in auditory processing and social interaction perception. In Experiment 1, monolingual participants listened to two-speaker conversations (intact or sentence-scrambled) and one-speaker narrations in both a known and unknown language. Speaker number and conversational coherence were explored in separately localised regions-of-interest (ROI). In Experiment 2, bilingual participants were scanned to explore the role of language comprehension. Combining univariate and multivariate analyses, we found initial evidence for a heteromodal response to social interactions in SI-pSTS. Specifically, right SI-pSTS preferred auditory interactions over control stimuli and represented information about both speaker number and interactive coherence. Bilateral temporal voice areas (TVA) showed a similar, but less specific, profile. Exploratory analyses identified another auditory-interaction sensitive area in anterior STS. Indeed, direct comparison suggests modality specific tuning, with SI-pSTS preferring visual information while aSTS prefers auditory information. Altogether, these results suggest that right SI-pSTS is a heteromodal region that represents information about social interactions in both visual and auditory domains. Future work is needed to clarify the roles of TVA and aSTS in auditory interaction perception and further probe right SI-pSTS interaction-selectivity using non-semantic prosodic cues.

**Highlights:**

- Novel work investigating social interaction perception in the auditory domain.
- Visually defined SI-pSTS shows a heteromodal response profile to interactions.
- Yet, it prefers visual to auditory stimuli. The reverse was found for anterior STS.
- Temporal voice areas show qualitatively different response compared to SI-pSTS.
- Future studies are needed to corroborate the unique role of right SI-pSTS.

## 1. Introduction

Every day, humans need to navigate social encounters which contain potent social cues that not only meet the eye (e.g., gestures and facial expressions) but also the ear (e.g., intonation and vocalisation timing). As such, gauging information from both visual and auditory social interactions is critical to support adaptive behaviour in a complex social world. Indeed, human sensitivity to interactions is such that the presence of an interaction facilitates processing speed (Papeo et al., 2019; Vestner et al., 2019), recognition accuracy (Papeo & Abassi, 2019; Papeo et al., 2017), and working memory efficiency (Ding et al., 2017; Vestner et al., 2019). Correspondingly, akin to evidence for face-(Kanwisher & Yovel, 2006), body-(Downing et al., 2001) or voice-selective (Belin et al., 2002; Belin et al., 2000) brain areas, neuroimaging studies have identified interaction-sensitive regions within bilateral lateral occipito-temporal cortex. Specifically, the posterior superior temporal sulcus social interaction region (SI-pSTS) has been found to play a key role in the representation of *visually* perceived dynamic social interactions. Across a range of stimuli that vary in the strength of relevant social cues (e.g., point-light displays, animated shapes, videos of dyads), the SI-pSTS responds about twice as strongly to interacting dyads compared to two independently acting individuals (Isik et al., 2017; Walbrin et al., 2018; Walbrin & Koldewyn, 2019) and shows this selectivity even in naturalistic videos (Landsiedel et al., 2022; Masson & Isik, 2021). Further, multivariate decoding analyses have found that the right SI-pSTS not only discriminates between interactors and non-interactors, but also appears to represent the type and emotional content of interactions (e.g., competing/cooperating; Isik et al., 2017; Walbrin et al., 2018; Walbrin & Koldewyn, 2019). On the other hand, extrastriate body area (EBA) has been implicated in the processing of the relational properties; i.e., the facing direction, between two bodies, which could be classed as ‘prototypical’ visual interactions (Papeo, 2020), for both static (Abassi & Papeo, 2020, 2021) and dynamic (Bellot et al., 2021) stimuli. Altogether, this suggests a special role of *visually* presented interactive cues within the social brain. However, although interactions in the world are conveyed through auditory as well as visual means, auditory cues to interaction have not received much attention thus far. The current study sought to address this.

Undeniably, perceiving interactions between others is not only a visual but also an *auditory* perceptual experience. For instance, a person might overhear two people behind them conversing with each other or listen to a radio interview. Even without visual information, a great deal can be derived about interactions not only based on *what* is heard, i.e., the semantic content, but also *how* that content is conveyed, i.e., the tone of interaction due to variations in prosody (e.g., changes in intonation or volume, use of pauses, etc.). Indeed, conversational characteristics can affect how interactors are perceived and which characteristics are attributed to them during conversations. For instance, when listening to a two-speaker conversation which culminates in one person asking for something, listener’s ratings of the respondent’s willingness to agree to this request decreased as the gap between the request and their affirmative response increased (Roberts & Francis, 2013; Roberts et al., 2006). Henetz (2017) used similar procedures and found that perception of a conversations’ awkwardness as well as interlocutors’ rapport and desire to interact in the future depended on inter-turn silences. This emphasizes that cues derived whilst listening to interactions are no less informative than cues gathered from visually observing interactions. In fact, one could suggest that most visual cues to interaction have auditory counterpoints that convey nearly identical social information including things like the identity of interactants (Awwad Shiekh Hasan et al., 2016; Stevenage et al., 2012), conversational turn-taking (Cañigueral & Hamilton, 2019; Pijper & Sanderman, 1994), as well as cues to the interactants’ emotions (de Gelder et al., 2015; Demenescu et al., 2014; Schirmer & Adolphs, 2017), intentions (Enrici et al., 2011; Hellbernd & Sammler, 2016), and social traits (Ponsot et al., 2018; Todorov et al., 2015).

Despite the richness of information contained in auditory interactions, the neural underpinnings of social interaction perception have been investigated almost exclusively in the visual domain. Studies probing neural representation of purely auditory interactions are next to non-existent. The closest proxy are studies investigating the auditory motion of two people walking, which convey some sense of togetherness or interactiveness. Bidet-Caulet et al. (2005) asked participants to listen to footsteps of two people walking, one on their left side (left ear) and one on their right side (right ear). Subsequently, one of the walkers would cross the auditory scene, thus their footstep sounds would move towards the same side as the other walker’s, which required the participants’ response. Compared to a simple noise detection task (requiring auditory attention), bilateral posterior superior temporal sulcus (pSTS) increased activation in the footstep condition. However, their study did not probe auditory interaction perception, per se, which would have required contrasting auditory motion of one person vs two. Work by Saarela and Hari (2008) tested this directly and found no differences in brain activation in pSTS, or indeed any other brain region, when comparing footsteps of two walkers vs one. While these auditory motion studies don’t provide evidence for a region that is selectively engaged by auditory interactions, to the best of our knowledge, no study has specifically investigated the perception of auditory interactions using actual conversational compared to non-conversational speech, or indeed probed whether regions characterised by its sensitivity to visual interactions might also be driven by auditory interactions.

In spite of the lack of interaction-specific studies, investigations focused on the processing of other social stimuli (e.g., faces and voices) support the notion of heteromodal (i.e. responding to both visual and auditory stimuli) processing in the broader STS region (Deen et al., 2020; Watson et al., 2014), which is in line with reports of a significant overlap between face- and voice-sensitive voxels in parts of the pSTS (Deen et al., 2015). Furthermore, several studies have proposed the STS as an area of audio-visual integration of both emotional and neutral facial and vocal expressions (Kreifelts et al., 2009; Robins et al., 2009; Watson et al., 2014; Wright et al., 2003), reflected by enhanced pSTS activation in response to both modalities compared to unimodal stimulus presentation. Given its proximity to auditory cortical areas, and nearby regions demonstrated to be integrative and/or heteromodal, the SI-pSTS seems an obvious candidate to investigate in the context of auditory interactions. This is in contrast to EBA, which may also be involved in interaction processing (Abassi & Papeo, 2020, 2021; Bellot et al., 2021), but which is considered to be a strictly visual region and not responsive to auditory information (Beer et al., 2013).

Across two experiments, we addressed the hypothesis that the SI-pSTS region might not only play a crucial role in the processing of visual but also auditory interactions using speech stimuli with three levels of interactiveness; interactions (conversations between two people) and their scrambled counterparts, as well as non-interactions (stories narrated by one person) in two languages. Importantly, scrambling recombined complete utterances taken from different interactions. Thus, speech was comprehensible at the sentence level but sentences were not semantically related to each other. In Experiment 1, participants were monolingual, whereas in Experiment 2, participants were bilingual. We used functional localisers (Fedorenko et al., 2010) to define bilateral regions of interest (ROIs); the visual SI-pSTS region, voice-selective temporal voice areas (TVA), and temporal parietal junction (TPJ) and tested their responses to auditory interactions across both univariate and multivariate pattern analyses. We hypothesized that if visual SI-pSTS was in fact heteromodal, it would show greater response to conversation stimuli involving two speakers compared to one-speaker narrations (regardless of semantic comprehension in monolinguals in Experiment 1). Beyond this broad test of auditory interaction sensitivity, we also expected SI-pSTS to be sensitive to the difference between conversations and scrambled conversations (which for monolingual participants deteriorated interactive cues of conversation coherence in their native language, and conversational flow/prosody in the unknown language). TVA (Agus et al., 2017; Belin et al., 2002; Belin et al., 2000) was included as an auditory control region that we expected to respond to all conditions, though given the dearth of information on auditory interaction processing, we did not have strong expectations regarding its interaction sensitivity. Our reasoning was that if SI-pSTS did not show heteromodal characteristics and sensitivity to auditory interactions could be ‘found’ anywhere in the brain, such sensitivity might emerge in an area tuned to voices, like TVA. Finally, TPJ was included as a ‘social’ control region that is spatially very near SI-pSTS but that we did not expect to either be driven by auditory stimuli in general, or manipulations of interactiveness specifically (Walbrin & Koldewyn, 2019; Walbrin et al., 2020). Additionally, whole-brain analyses were conducted to explore the wider brain networks implicated in auditory interaction processing.

## 2. Methods

### 2.1 Experiment 1

#### 2.1.1 Participants

Twenty-four right-handed participants were recruited to take part in this study. Handedness was confirmed using the Edinburgh Handedness Inventory (EHI, Oldfield, 1971). All participants had normal or corrected to normal vision, were native English speakers, and had no German language skills. After data exclusion (see 2.1.2), one participant was removed from the analyses (final sample of N = 23; mean age = 22.35, SD = 3.04; 7 males). All participants gave informed consent, were debriefed at the end of the study, and received monetary renumeration for their time. The protocol was approved by the School of Psychology’s ethics committee at Bangor University and was pre-registered on AsPredicted.org (ID23865) on 23/05/2019.

#### 2.1.2 Design & Procedure

To investigate auditory interaction perception with and without language comprehension, the main experimental task consisted of a 2 × 3 repeated measures fMRI event-related design. Across two *languages* (English and German), auditory interactiveness was manipulated using three *conditions*: non-interactive one-speaker narrations, interactive two-speaker conversations, as well as an intermediate condition using scrambled conversations (see 2.1.3 for details). This condition still contained interactive cues (two speakers taking turns); however, conversational content was not coherent. German stimuli were used to explore interactive effects independent of stimulus comprehension.

Each run contained 36 trials, including 24 task trials (4 per condition) and 12 catch trials (2 per condition). The order of conditions was pseudo-randomised using custom MATLAB code to optimise the efficiency of the design. The inter-stimulus interval was jittered (mean jitter 1.5s, range =0s – 3s). Participants completed seven runs (28 task trials and 14 catch trials per condition across runs, thus 252 trials overall), each between 5.6 – 5.8 minutes in length. Due to the variability in stimulus length (range: 6 -11 seconds), each run contained a specific set of stimuli. The order of the runs was counter-balanced across participants.

Participants were instructed to listen attentively, and an orthogonal catch-trial detection task was used to maintain (and confirm) participants’ attention throughout. Catch trials consisted of recordings that were manipulated such that a single word was repeated, e.g., “Do you *need need* anything from the supermarket?”. The occurrence of these repeated catch words was balanced for all conditions such that the repeated word occurred equally often in the first, second, third or fourth sentence of a stimulus. Participants had to accurately detect seven out of 12 catch trials per run for that run to be included in subsequent analyses (see Supplementary S1 for the behavioural results). Based on this criterium, two participants had one, and another participant three run(s) removed from the analyses. Only participants with five or more runs after data exclusion were included in the final sample.

The task was presented in Psychtoolbox 3.0.14 (Brainard, 1997; Kleiner et al., 2007) using MATLAB 2018.a (The MathWorks Inc.) running on a Linux Ubuntu 16.04 distribution stimulus computer. Sensimetrics (model S15) MR-safe in-ear earphones were used for stimulus presentation. Stimulus volume was adjusted to a comfortable level individually for each participant.

#### 2.1.3 Stimuli

Scripted conversations and narrations were developed in English, and subsequently translated into German by two native speakers. All stimuli were recorded specifically for this study (see Supplementary S2 for recording details) by native English or native German speakers. A large set of stimuli was recorded, from which the final stimulus set was selected (see Supplementary S3 and S4.1 for details).

*Narrations* were recorded for each speaker separately. Narration content was loosely based on and inspired by children’s books. Care was taken that narrations remained descriptive rather than invoking mentalizing processes.

*Conversations* were recorded in pairs (two same-gender pairs per language condition, one male, one female) to capture “true” interactions. They consisted of a short exchange (usually four sentences in total) between the two speakers taking turns (Agent A - Agent B - Agent A - Agent B). All conversations were recorded twice so that both speakers played both agent roles. Conversations varied in content, e.g., asking a friend about their exam or ordering food in a restaurant.

*Scrambled conversations* were created from the original conversation scripts by randomly combining individual speaker turns from different conversations into a new combination, which still consisted of two speakers taking turns, but where the turns were taken from different conversations and, thus, were unrelated in semantic meaning. Importantly, this process could result in a speaker turn being spoken by a different speaker in the scrambled compared to the original conversations whilst the script; i.e., the semantic content remained the same. For this, the original conversations (from each gender pair) were cut up into their individual speaker turns (two per agent) using Audacity (The Audacity Team) audio software. Using custom MATLAB code, turns from four random original conversations were selected and re-combined such that each turn was no longer in its original position of the conversation. For example, the opening turn of a conversation could only appear in the 2^nd^, 3^rd^, or 4^th^ position in a scrambled conversation. This method was chosen to disrupt the natural conversational flow not only through mixing up content but also disrupting prosodic and intonational cues. Scrambling generated equal numbers of stimuli with either speaker taking the role of agent A or agent B.

All stimuli were analysed in Praat software (http://www.praat.org) to assess mean pitch (fundamental frequency, F0) for each condition (see Supplementary S4.2). Furthermore, the final set of English stimuli was rated on perceived naturalness, valence, imaginability/mental imagery, and for the two-speaker conditions also on interactiveness and perceived closeness between the two speakers. In brief, conversation and narrations were closely matched on all non-interactive dimensions. In contrast, conversations and scrambled conversations significantly differed across all scales (see Supplementary S4.3 for rating data and statistics).

#### 2.1.4 Localiser tasks

To define independent regions of interest (ROI), participants also completed a set of established localiser tasks. The *interaction localiser* (Isik et al., 2017; Walbrin et al., 2018) was used to localise the SI-pSTS region sensitive to visual social interactions. Across three runs, participants watched videos of two point-light agents in three conditions (interaction; scrambled interactions, and non-interaction/ independent actions). Each run contained two 16-second blocks per condition and three 16-second rest blocks (total run time 144 seconds). Bilateral SI-pSTS was localised using the contrast interactions >. non-interactions. The *voice localiser* (Pernet et al., 2015) was used to localise the temporal voice areas (TVAs) along the anterior-posterior axis of the STS. The TVAs show sensitivity to human vocal (speech and non-speech) sounds (Agus et al., 2017; Belin et al., 2002; Belin et al., 2000; Pernet et al., 2015), and seem particularly involved in processing of paralinguistic information such as gender (Charest et al., 2013), identity (Latinus et al., 2013), or emotion (Ethofer et al., 2012; Ethofer et al., 2009). Therefore, we do not expect to find interaction-sensitivity in the TVAs. Participants completed one run, listening to human vocal sounds (speech, e.g., words, or syllables; and non-speech sounds, e.g., laughs or sighs); and non-vocal sounds (natural sounds like waves or animals, and man-made object sounds like cars or alarms) across twenty 8-second blocks respectively. These condition blocks were interspersed with twenty 10-second blocks of silence (total run time 10.3-minutes). Bilateral TVA was localised using the contrast human vocal sounds > non-vocal sounds. Finally, a third localiser was used to define temporo-parietal junction as a control region within the ‘social brain’ (see Supplementary S5 for details).

#### 2.1.5 MRI parameters, pre-processing & GLM estimation

Data was collected at the Bangor Imaging Centre using a Philips Achieva 3-T scanner using a 32-channel head coil (Philips, Eindhoven, the Netherlands). A T2*-weighted gradient-echo single-shot EPI pulse sequence (with SoftTone noise reduction, TR = 2000ms, TE = 30ms) was used for all tasks (with slightly different parameters depending on task for flip angle, FOV, number of slices and slice order, see Table 1).

**Table 1.**
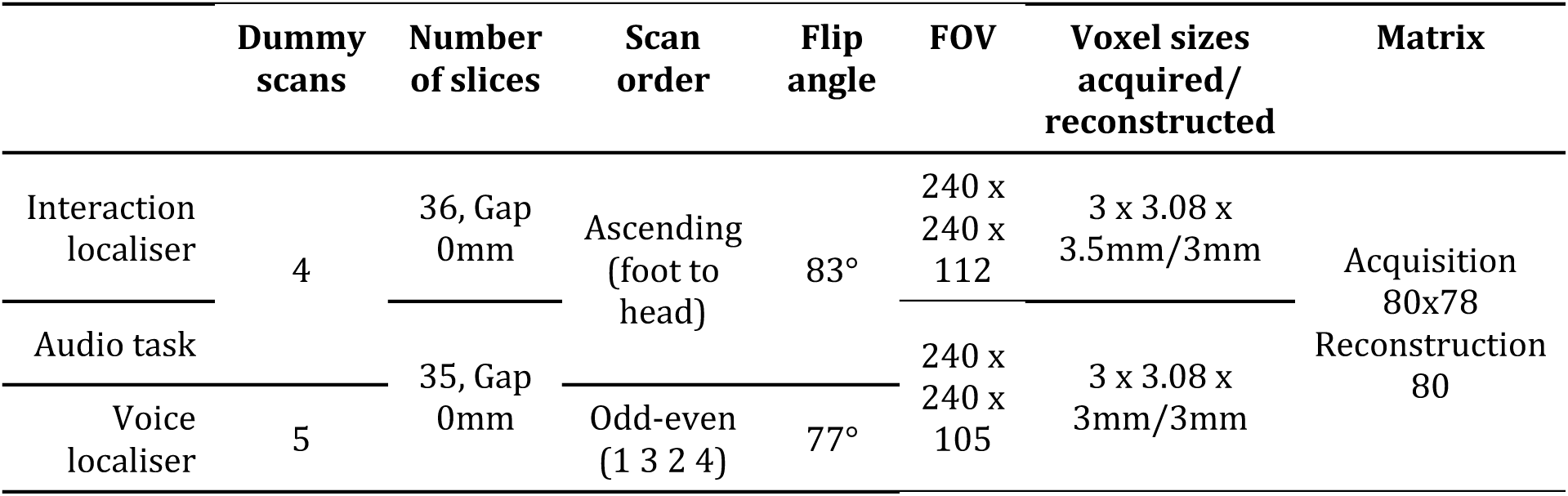
Scanning parameters

Structural images were obtained with the following parameters: T1-weighted image acquisition using a gradient echo, multi-shot turbo field echo pulse sequence, with a five echo average; TR= 12 ms, average TE = 3.4 ms, in 1.7 ms steps, total acquisition time = 136 s, flip angle = 8°, FOV = 240 × 240, acquisition matrix = 240 × 224 (reconstruction matrix = 240); 128 contiguous axial slices, acquired voxel size(mm) = 1.0 × 1.07 × 2.0 (reconstructed voxel size = 1mm^3^).

Pre-processing and general linear model (GLM) estimation were performed using SPM 12 (fil.ion.ucl.ac.uk/spm/software/spm12) in MATLAB 2018.a (The MathWorks Inc.). Pre-processing included slice-timing (event-related main auditory task only), realignment & re-slicing, co-registration to anatomical image, segmentation, normalization (normalised MNI space with 2 mm isotropic voxels), and smoothing. All SPM12 default pre-processing parameters were used except for the use of an initial 3mm FWHM Gaussian smoothing kernel. This smoothing kernel is recommended when using ArtRepair toolbox (v5b, Mazaika et al., 2005). Specifically, ArtRepair was used to detect and repair noisy volumes (volumes that contained more than 1.3% variation in global intensity or 0.5mm/TR scan-to-scan-motion). 13 subjects required repairs in at least one run. Prior to first-level modelling, data was smoothed again using a 5mm FWHM kernel. Subsequently for each run, event or block durations and onsets for each experimental condition were modelled using a boxcar reference vector and convolved with a canonical hemodynamic response function (without time or dispersion derivatives) using a 128-second high-pass filter and autoregressive AR(1) model. Head motion was modelled using six nuisance regressors (translation and rotation). Additionally, for the main auditory task, catch trials were modelled as a regressor of no interest. Rest periods were modelled implicitly.

#### 2.1.6 Whole-brain group analyses

For localiser tasks, respective contrasts were modelled at the group level using one-sample t-tests. For the main auditory task, the Multivariate and Repeated Measures (MRM) toolbox (McFarquhar et al., 2016) was used. Each participant’s baseline contrast images of the six experimental conditions were entered into a 2 (Language) × 3 (Condition) repeated-measures ANOVA. Within this model, *F*-contrasts were computed for the main effects and the interaction effect using Wilks’ Lambda as multivariate test statistic. All reported F-contrasts were thresholded using an initial cluster-forming threshold of *p* < 0.001 uncorrected, followed by permutation tests (5000 iterations) to provide cluster-level FWE correction for multiple comparisons of *p_FWE_* < 0.05.

#### 2.1.7 ROI creation & percent signal change (PSC) analyses

Functional ROIs were defined following a ‘group-constrained’ ROI definition approach (for details see Julian et al. (2012)). This ensured that ROI definition was as objective as possible. To start, group-level T/F-maps were used to identify MNI coordinates of bilateral ROIs (see Supplementary S6). These coordinates formed the centres of initial *8mm bounding spheres*. Subject-specific search spaces were then defined by running a group-level analysis to determine a peak coordinate for activation that was used to localise this search space using a leave-one-subject-out (LOSO) approach, i.e., the group contained all subjects except the ‘current’ subject whose search space was being defined. The final *subject-specific search space* was defined based on the intersection of the original 8mm bounding sphere with the group-level T-map of the LOSO scheme. To create the final *subject-specific ROI* (see Figure S1), the top 100 *contiguous* voxels (highest T-values) within the *subject-specific search* were selected for each participant individually (Walbrin et al., 2020). Thus, while all ROIs were 100 voxels in size, they differed across participants in their exact placement. Additionally, a leave-one-run-out (LORO) scheme was implemented in this step (defining an ROI on all but one run, extracting PSC from the remaining one in an iterative n-fold partition scheme) in cases where ROI definition and extraction were based on the same task, i.e., when testing the response of the SI-pSTS itself to the interaction localiser. This procedure could not be applied when testing the response of the TVA to the voice localiser conditions, however, as the localiser consisted of only one run. PSC data were extracted from ROIs using the Marsbar toolbox (Brett et al., 2002).

For the main auditory task, PSC were analysed in a 2 (Language) × 3 (Condition) repeated-measures ANOVAs for each ROI respectively. Greenhouse-Geisser correction for violation of assumptions of sphericity were applied where necessary. Multiple comparison correction was implemented based on the number of ROIs tested in a given contrast; i.e. multiple tests for each contrast were considered as a “family” of statistical tests that should be corrected across. Given our four regions of interest (bilateral SI-pSTS and TVA), this resulted in a corrected *p*-value of *p* < .0125 (0.05/4) for both contrasts used to test main and interaction effects. For the interaction localiser, differences between conditions were analysed using paired-sample t-tests, with particular focus on two contrasts of interest: interactions vs. non-interactions and scrambled interactions respectively. For the auditory task, multiple comparison correction was applied based on the number of ROIs tested in a given contrast (corrected p-value of *p* < .0125). Furthermore, for selected ROIs, auditory- and visual-*interaction selectivity* was calculated as the t-value of differences in activation between conversations vs narrations across languages for the main experimental task, and interactions vs non-interactions for the interaction localiser, and compared using paired-sample t-tests.

For all paired t-test comparisons, effect sizes represent Cohen′s d for repeated measures (*d_rm_*) which represents the mean difference standardized by the standard deviation of the difference scores corrected for the correlation between the measurements (Lakens, 2013).

#### 2.1.8 Multivariate pattern analyses (MVPA)

Pattern decoding analysis using an iterative n-folds partition scheme of LORO was implemented using the CoSMoMVPA toolbox (Oosterhof et al., 2016) with a focus on four contrasts of interest: conversations vs narrations and conversations vs scrambled conversations for each language. For each subject, a binary linear support vector machine (SVM) classifier was trained on a given ROI’s voxel patterns for the conditions of interest (i.e. beta values in a subject’s respective top 100-voxels ROI, see section 2.1.7, averaged across all trials per condition per run) in all but one run of data–with the ‘left-out’ run of data used to independently test classification performance. This resulted in as many folds of cross-validation as a subject had valid task runs (usually seven). Prior to classification, voxels patterns were normalized (demeaned) for each run separately. Following cross-validation, classification accuracy was averaged across all n-folds iterations before being entered into group level analysis. For each contrast of interest, average classification accuracy was tested against chance (50%) using one-tailed one-sample t-tests.

To correct for multiple comparisons across four ROIs, we used a so-called “singleton” neighbourhood, where each ROI to be corrected for was treated as one feature. This means that each ROI was only a neighbour of itself. This neighbourhood was then tested using monte-carlo based clustering statistics. Here, we used Threshold-Free-Cluster-Enhancement (TFCE, Smith & Nichols, 2009) as a clustering statistic with 10.000 iterations of Monte Carlo simulations (cf. cosmo_montecarlo_cluster_stat.m). Although traditionally, TFCE is used to test cluster survival based on iteratively testing the spatial clustering at different height thresholds to determine how much local support each feature (voxel) has (using a neighbourhood composed of many features), the same principle can also be applied to correct for multiple comparison across ROIs when conceptualising each ROI as a cluster (cf. cosmo_singleton_neighborhood.m). Each iteration of Monte Carlo simulations generated null data based on the sample’s classification accuracies in each ROI using a sign-based permutation approach (also implemented in FieldTrip; see Maris and Oostenveld, 2007). Significance was determined based on the comparison of “clustering” in the null data across all iterations compared to “clustering” observed in the original data. This method yields conservative estimates of significance. We report the resulting one-tailed Z-scores and p-values where Z-scores greater than 1.65 are indicative of significant above chance classification in each ROI after correction for multiple comparisons.

### 2.2 Experiment 2

#### 2.2.1 Participants

Twelve German native speakers took part in this study (mean age = 22.50, SD = 2.28, 3males). All participants were right-handed as confirmed by the EHI, and had normal or corrected to normal vision. The sample’s English language skills were at a minimum of B2 (upper intermediate, Common European Framework of Reference for Languages), which is the minimum level required by universities for international first year students. Participants gave informed consent and were debriefed and paid at the end of the study.

#### 2.2.2 Design & Procedure

All procedures were the same as in Experiment 1 although participants only completed the main auditory experimental task as well as the pSTS interaction localiser. Due to technical difficulties, the TVA voice localiser could not be acquired for this sample.

#### 2.2.3 MRI parameters & pre-processing

Compared to Experiment 1, images were acquired on a different scanner (Philips Igenia Elition X 3T scanner) with a 32-channel head coil (Philips, Eindhoven, the Netherlands) at the Bangor Imaging Centre. Acquisition parameters for functional runs were the same as in Experiment 1. The structural sequence was slightly different: for each participant, a high-resolution anatomical T1-weighted image acquired using a gradient echo, multi-shot turbo field echo pulse sequence, with a five-echo average; TR = 18ms, average TE = 9.8ms, in 3.2ms steps, total acquisition time = 338s, flip angle = 8°, FOV = 224x224, acquisition matrix = 224x220 (reconstruction matrix = 240); 175 contiguous slices, acquired voxel size (mm) = 1.0x1.0x2.0 (reconstructed voxel size = 1mm³).

Pre-processing and GLM estimation were performed using the same pipeline as in Experiment 1. Due to low levels of head-motion in this sample, ArtRepair was not used to repair noisy volumes. Due to human error, the first functional run of the main auditory task had to be discarded for two participants.

#### 2.2.4 ROI creation & PSC analyses

For the SI-pSTS, the same group constrained ROI definition process was used as in Experiment 1, resulting in subject-specific ROIs consisting of the top 100 contiguous voxels in each hemisphere separately. However, due to the small sample size in this study, the initial 8mm constraining sphere used the same group-level MNI coordinates as in Experiment 1, rather than using coordinates of our underpowered sample. Due to the missing voice localiser scan, for bilateral TVA, ROIs consisted of 6mm spheres, again using centre MNI coordinates from Experiment 1. This radius was chosen to select a sphere size that contained a comparable number of voxels as the SI-pSTS ROI (6mm sphere = 123 voxels). As before, PSC data were extracted for the experimental conditions of the main auditory task using Marsbar toolbox for the respective ROIs and analysed using 2 × 3 repeated-measures ANOVAs.

## 3. Results

### 3.1 Experiment 1

Results of the PSC results (Figure 1 left panel) and MVPA analyses (Figure 1 right panel) will be presented below separately for the visually defined SI-pSTS and auditorily defined TVA (refer to Table 2 for ANOVA statistics and Supplementary S7 Table S7 for condition means). Results for TPJ as an additional control region within the ‘social brain’ can be found in Supplementary S6, as it consistently showed activation at or below baseline. Bspmview toolbox was used for whole-brain data visualisation (DOI: 10.5281/zenodo.168074, see also https://www.bobspunt.com/software/bspmview/).

**Figure 1.**
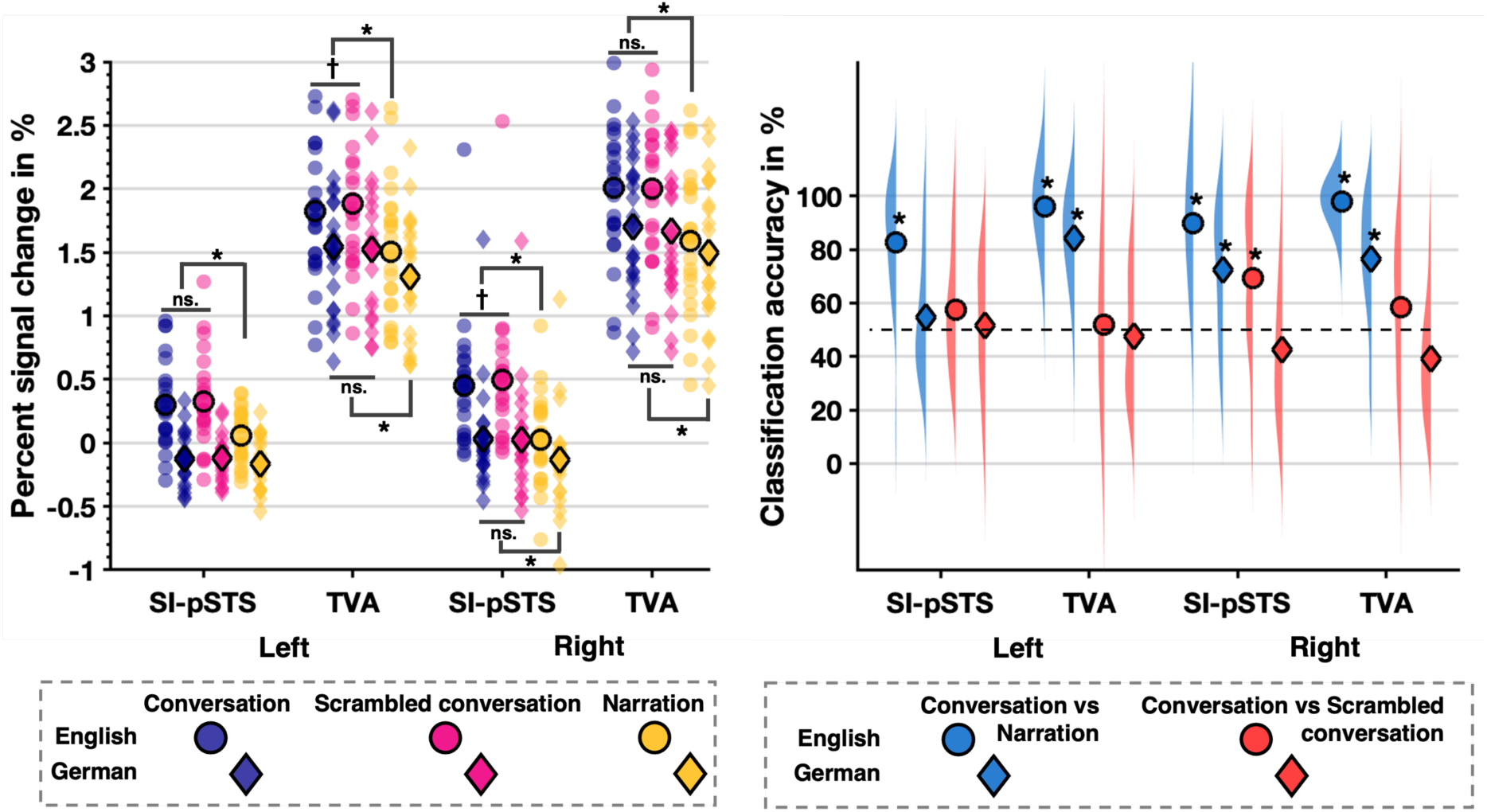
Condition means (circle/diamond shape with bold edge) and data distribution (scatter or density plot) for each ROI and hemisphere for Experiment 1. **Left panel**: Percent signal change data. Only significant post hoc t-test results are marked (*: p < .0125 corrected, †: p < .05 uncorrected). Effects for English conditions are shown above, and for German conditions below the condition mean. **Right panel**: SVM classification accuracy. Significant above chance classification accuracy is indicated using an asterisk (*: p < .05 TFCE-corrected). Chance level of 50% is represented using the dashed horizontal line.

**Table 2.** ANOVA results of PSC analyses for main auditory task in Experiment 1 and 2

|  |  | Experiment 1 |  |  |  | Experiment 2 |  |  |  |
| --- | --- | --- | --- | --- | --- | --- | --- | --- | --- |
|  |  | SI-pSTS |  | TVA |  | SI-pSTS |  | TVA |  |
|  |  | L | R | L | R | L | R | L | R |
| Main effect of Language | *F* | 36.02 | 50.58 | 29.44 | 15.84 | 1.37 | 1.71 | 4.91 | 13.56 |
|  | *p* | <.001 | <.001 | <.001 | <.001 | .27 | .22 | .049 | .004 |
| | $\eta_p^2$ | .62 | 0.70 | 0.57 | 0.42 | 0.11 | 0.13 | 0.31 | 0.55 |
| Main effect of Condition | *F* | 14.24 | 35.99 | 62.21 | 40.89 | 10.57 | 23.88 | 27.47 | 20.2 |
|  | *p* | <.001 | <.001 | <.001 | <.001 | <.001 | <.001 | <.001 | <.001 |
| | $\eta_p^2$ | 0.39 | 0.62 | 0.74 | 0.65 | 0.49 | 0.69 | 0.71 | 0.65 |
| Interaction effect | *F* | 12.18 | 24.71 | 9.86 | 25.42 | 0.81 | 3.5 | 0.10 | 0.25 |
|  | *p* | <.001 | <.001 | <.001 | <.001 | .41 | .048 | .91 | .78 |
| | $\eta_p^2$ | 0.36 | 0.53 | 0.31 | 0.54 | 0.07 | 0.24 | 0.01 | 0.02 |

#### 3.1.1 How does the visually defined SI-pSTS respond to auditory interactions?

##### PSC analyses

In line with our predictions, there was a main effect of Condition, where both conversations (rSI-pSTS: *t*(22) = 6.17, *p* < .001, *d_rm_* = 0.41 , lSI-pSTS: *t*(22) = 3.82, *p* < .001, *d_rm_* = 0.22) and scrambled conversations (rSI-pSTS: *t*(22) = 6.13, *p* < .001, *d_rm_* = 0.44, lSI-pSTS: *t*(22) = 4.00, *p* < .001, *d_rm_* = 0.20) evoked greater responses than narrations in bilateral SI-pSTS. However, PSC did not differ for intact vs. scrambled conversations. Thus, this effect was driven entirely by a difference between hearing two speakers vs. hearing only one. Unexpectedly, there was also a large main effect of Language, where responses in bilateral SI-pSTS were greater for English compared to German stimuli. Indeed, while PSC in bilateral SI-pSTS was significantly above baseline for English conversations and English scrambled conversations (all *t*(22) ≥ 4.06, all *p*s < .001), for lSI-pSTS, German conditions led to a significant decrease in activation (all *t*s(22) < -2.95, all *p*s < .007) and response in rSI-pSTS was not significantly different than baseline. Both effects suggest that the *comprehensibility* of heard interactions is important to response within the SI-pSTS. These main effects were qualified by a significant Language × Condition interaction. Whilst responses in lSI-pSTS were only greater for the comprehensible English two-speaker conversation compared to single-speaker narrations (*t*(22) = 4.14, *p* < .001, *d_rm_* = 0.77), rSI-pSTS showed this pattern independent of comprehensibility (English: *t*(22) = 6.28, *p* < .001, *d_rm_* = 0.90; German: *t*(22) = 4.76, *p* < .001, *d_rm_* = 0.41). Additionally, rSI-pSTS response to English scrambled conversations was slightly greater than conversations, albeit at an uncorrected level only (*t*(22) = -2.16, *p* = .04, *d_rm_* = -0.09), suggesting weak sensitivity to the *coherence* of comprehensible interactions.

##### MVPA analyses

Classification analyses in the rSI-pSTS revealed that the SVM classifier could discriminate between voxel patterns representing conversations and narrations in both languages (English: M = 0.90, SE = 0.03, Z = 3.72, *p_TFCE_* < .001; German: M = 0.72, SE = 0.05, Z = 3.24, *p_TFCE_* < .001), in line with the PSC results. Crucially, strengthening the PSC results, the classifier could also decode voxel-patterns of English conversations vs. scrambled conversations (M = 0.69, SE = 0.06, Z = 2.49, *p_TFCE_* = .006) with above chance accuracy. This suggests that rSI-pSTS voxel-patterns code for interaction information based on the number of speakers and interaction coherence when two speakers are present. Classification analyses in lSI-pSTS revealed above chance discrimination between English conversations vs narrations only (M = 0.83, SE = 0.04, Z = 3.72, *p_TFCE_* < .001), suggesting no strong auditory interaction sensitivity in the left hemisphere. German conversations vs. German scrambled conversations were not decodable above chance in either region.

#### 3.1.2 How does the TVA, a region generally sensitive to voices, respond to auditory interactions?

##### PSC analyses

In bilateral TVA, while there was also a significant main effect of Language, PSC was significantly greater than baseline for all conditions, regardless of language (all *t*(22) ≥ 12.62, all *p*s < .001). Thus, bilateral TVA was clearly driven by voice stimuli regardless of comprehensibility. However, bilaterally, PSC was greater in response to English compared to German stimuli. As in the SI-pSTS, there was also a main effect of Condition bilaterally. PSC was smaller for narrations compared to both conversations (rTVA: *t*(22) = 7.10, *p* < .001, *d_rm_* = 0.54, lTVA: *t*(22) = 7.28, *p* < .001, *d_rm_* = 0.54) and scrambled conversations (rTVA: *t*(22) = 6.09, *p* < .001, *d_rm_* = 0.55, lTVA: *t*(22) = 9.58, *p* < .001, *d_rm_* = 0.51) but no difference was found between the latter two. Thus, the number of voices, hearing one or two speakers, clearly modulated TVA activation. These main effects were qualified by a significant Language × Condition interaction. Bilaterally, TVA responses were greater for conversations compared to narrations for both English (rTVA: *t*(22) = 7.57, *p* < .001, *d_rm_* = 0.74, lTVA: *t*(22) = 5.66, *p* < .001, *d_rm_* = 0.55) and German stimuli (rTVA: *t*(22) = 5.18, *p* < .001, *d_rm_* = 0.36, lTVA: *t*(22) = 8.98, *p* < .001, *d_rm_* = 0.44). Surprisingly, although at an uncorrected level, lTVA responded less to English conversations compared to scrambled conversations (*t*(22) = -2.62, *p* = .02, *d_rm_* = -0.09), indicating potential sensitivity to the *coherence* of comprehensible interactions.

##### MVPA analyses

Classification analyses revealed that voxel patterns representing conversations and narrations could be decoded for each language respectively in right (English: M = 0.98, SE = 0.01, Z = 3.72, *p_TFCE_* < .001; German: M = 0.77, SE = 0.04, Z = 3.54, *p_TFCE_* < .001), and left (English: M = 0.96, SE = 0.02, Z = 3.72, *p_TFCE_* < .001; German: M = 0.84, SE = 0.03, Z = 3.72, *p_TFCE_* < .001) TVA (consistent with the PSC results). Importantly, bilaterally, discrimination of conversations vs. scrambled conversations based on TVA voxel patterns was not successful for either language.

#### 3.1.3 What is the evidence for heteromodal social interaction processing in SI-pSTS (and TVA)?

To examine heteromodal processing in response to social interactions, we examined and, if appropriate, compared responses to visual interactions with responses to auditory interaction within our ROIs. Analyses confirmed sensitivity to visual interactions in bilateral SI-pSTS , which showed greater responses to interactions compared with both non-interactions and scrambled interactions (all *t*s(22) > 5.84, all *p*s < .001; see Figure 2 left panel and Supplementary S7, Table S8). In contrast, bilateral TVA responded at or below baseline to the interaction localiser conditions (all *t*(23) < -2.00, *p* < .06). Therefore, the subsequent comparison of interaction-selectivity across modalities focused on SI-pSTS only. Analyses revealed that visual interaction selectivity (rSI-pSTS: M = 0.62, SE = 0.08, lSI-pSTS: M = 0.53, SE = 0.06) in bilateral SI-pSTS was significantly greater (rSI-STS: *t*(22) = 5.81, *p* < .001, *d_rm_* = 0.34, lSI-pSTS: *t*(22) = 6.60, *p* < .001, *d_rm_* = 0.26) than auditory interaction selectivity (rSI-pSTS: M = 0.29, SE = 0.05, lSI-pSTS: M = 0.14, SE = 0.04). Altogether, this suggests heteromodal processing of social interaction in SI-pSTS, though with clear preference for visual stimuli.

**Figure 2.**
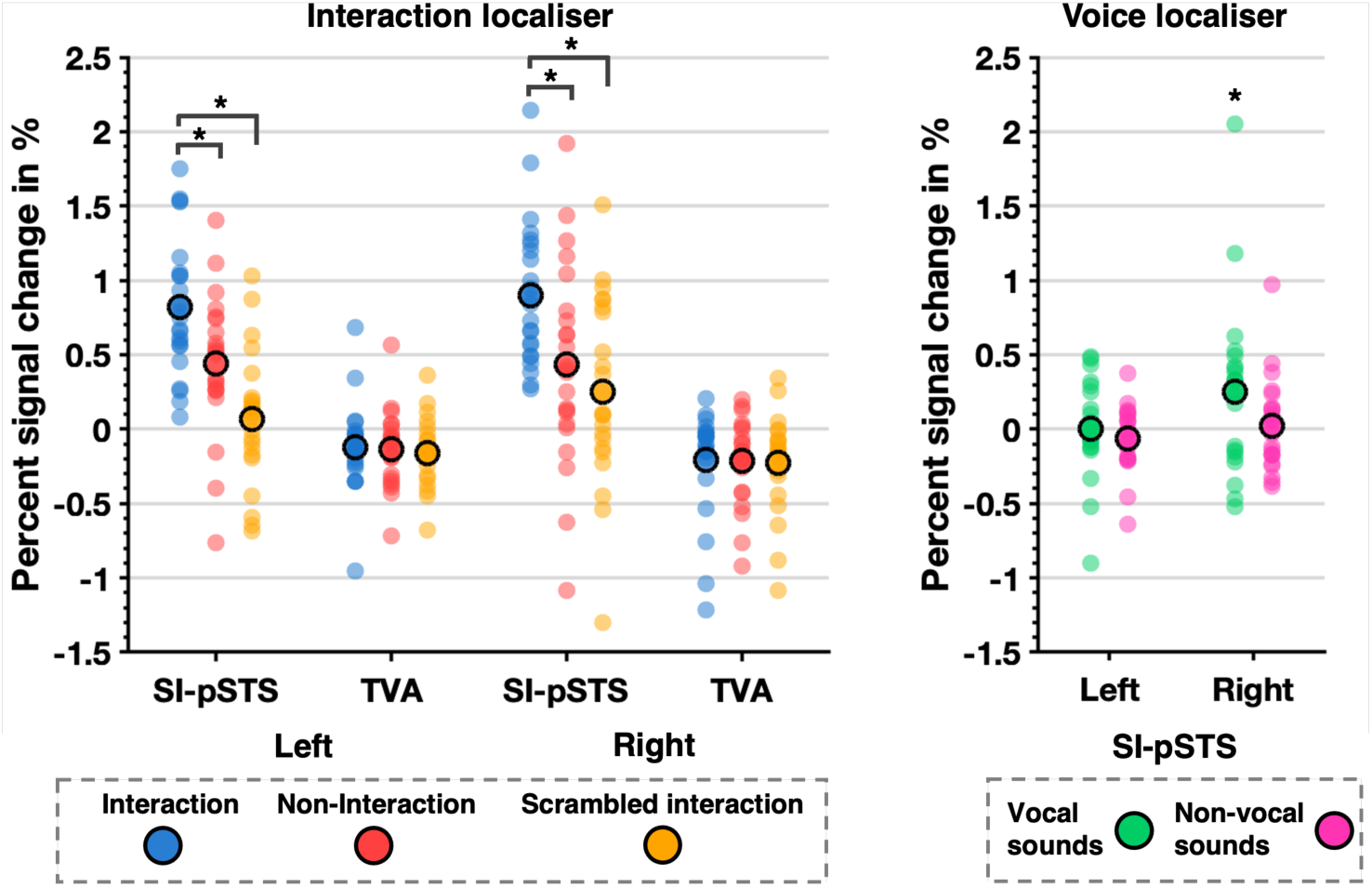
Illustration of PSC condition means (circle with bold edge) and data distribution (scatter plot) for localiser data of Experiment 1. **Left panel:** Interaction localiser data for SI-pSTS and TVA. Significant condition differences are marked (* p < .0125 corrected). **Right panel:** Voice localiser data for SI-pSTS only. Above baseline responses are marked (* p < .05).

#### 3.1.4 Does general voice sensitivity explain responses to auditory interactions in SI-pSTS?

General responsiveness to voice stimuli (see also Supplementary S7, Table S9) was examined by extracting PSC from the voice localiser in SI-pSTS (see Figure 2, right panel). Although vocal sounds activated rSI-pSTS above baseline (*t*(23) = 2.16, *p* = .04), the region was not strongly driven by human voices. In fact, compared to comprehensible English conversations (M = 0.45, SE = 0.10), rSI-pSTS responded about 50% less to vocal stimuli (M = 0.25, SE = 0.12). Overall, this analysis suggest, that SI-pSTS responses in the main experimental task were *not* due to a general sensitivity to voices.

#### 3.1.5 Does whole-brain data reveal an additional region sensitive to auditory interactions?

To explore whole-brain auditory interaction sensitivity, we followed a data-driven approach. Rather than focussing on the main effect of condition, the ROI PSC indicated an unexpected but robust Language × Condition interaction effect for our key ROIs. Therefore, the whole-brain interaction effect contrast was used to identify potential candidate regions that may be sensitive to auditory interactions. As it is evident from both Figure 3 and Table 3, brain activity was modulated by our factors within large clusters in bilateral STS, including substantial portions of the sulcus along much of its anterior-posterior axis. Other activations included prefrontal clusters in bilateral inferior frontal gyrus, right middle frontal gyrus, left superior medial gyrus, right anterior cingulate cortex, left precuneus, as well as the right cerebellum (Crus 2). Bilaterally, the global peak of *F*-values fell within the anterior portion of the STS clusters. In an exploratory post-hoc analysis, coordinates close to this peak were used to define bilateral anterior STS (aSTS) ROIs for PSC extraction using an iterative LORO process (see 2.1.7 above).

**Figure 3.**
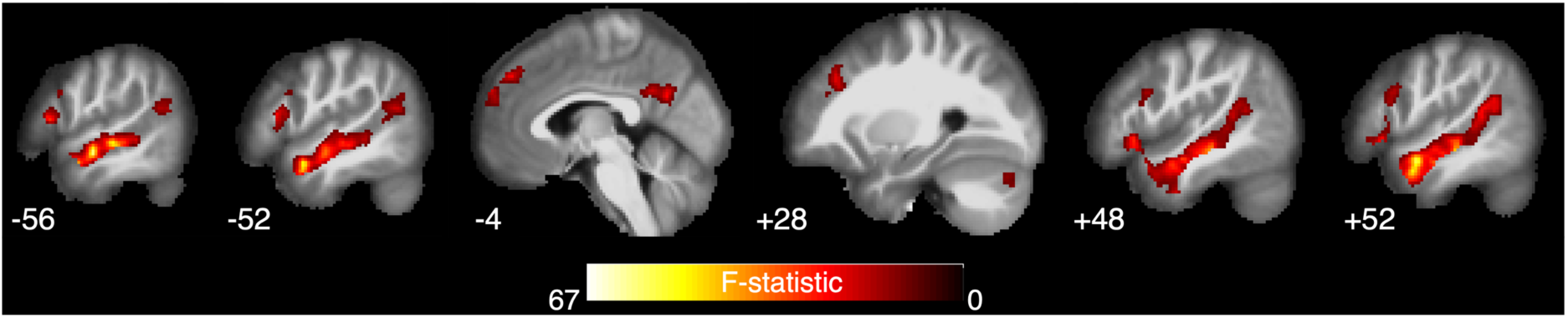
Sagittal view of whole-brain group analysis Language × Condition interaction F-contrast. Slices in MNI space with x-coordinate shown next to each slice.

**Table 3.**
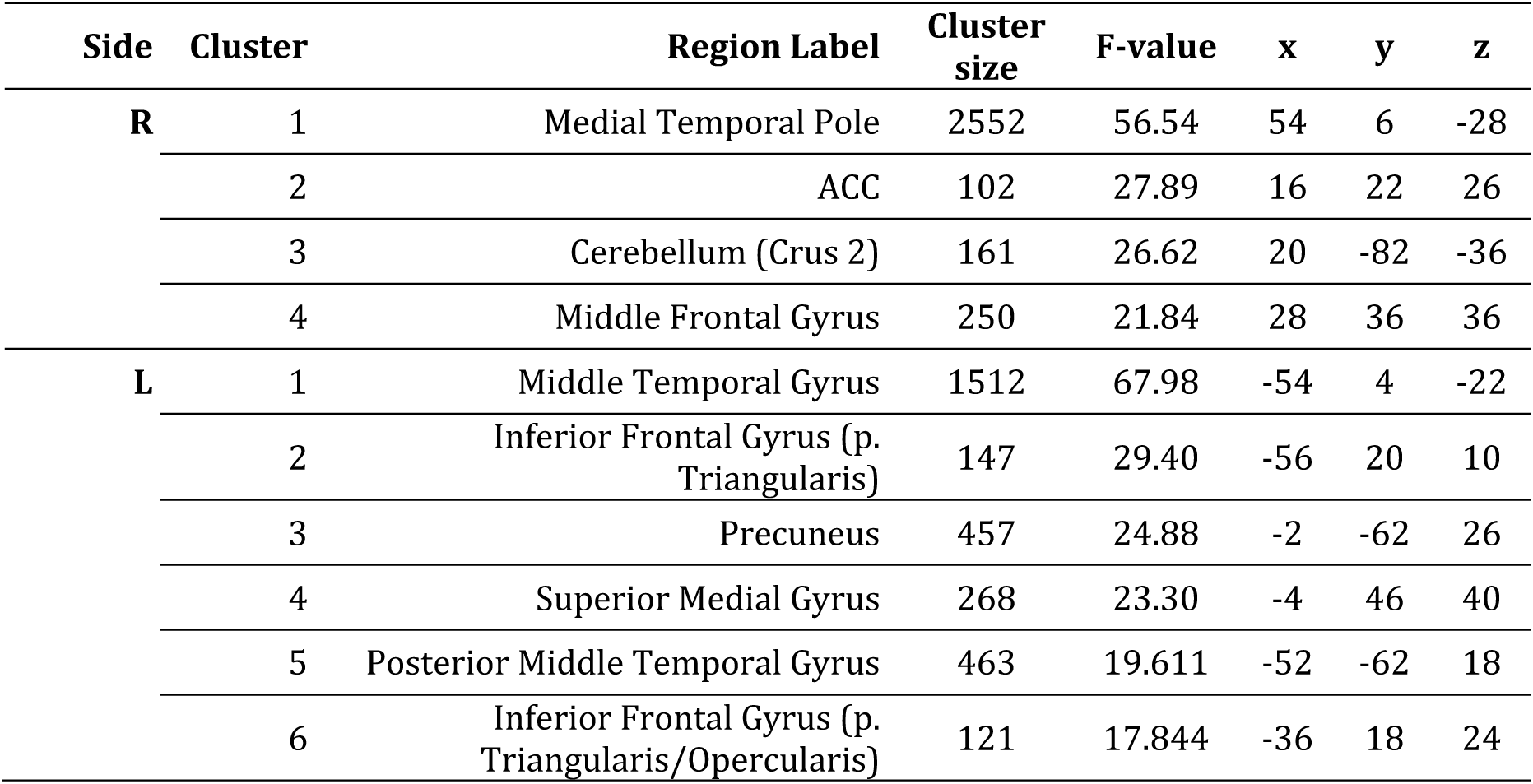
Significant clusters for whole-brain Language × Condition interaction F-contrast, cluster-corrected p_FWE_ < 0.05. All x, y, and z coordinates in MNI space.

As in both SI-pSTS and TVA regions, PSC analyses revealed a significant main effect of Condition in bilateral aSTS, indicating sensitivity to auditory interactions (see Table 4 Table 4and Figure 4 left panel, as well as Supplementary S7, Table S10 for condition means). PSC was smaller for narrations (compared to both conversations (raSTS: *t*(22) = 9.69, *p* < .001, *d_rm_* = 0.22, laSTS: *t*(22) = 7.29, *p* < .001, *d_rm_* = 0.15) and scrambled conversations (raSTS: *t*(22) = 8.60, *p* < .001, *d_rm_* = 0.22, laSTS: *t*(22) = 8.85, *p* < .001, *d_rm_* = 0.13). Interestingly, right but not left aSTS responded more to conversations compared to scrambled conversations (*t*(22) = 3.08, *p* = .005, *d_rm_* = 0.24), indicating that the region was not merely driven by the difference of hearing two speakers vs hearing one. Furthermore, there was a main effect of Language. Responses were greater for English compared to German stimuli. Indeed, PSC in bilateral aSTS was significantly above baseline for English conversations and English scrambled conversations, and for left aSTS also for English narrations (all *t*s(22) ≥ 6.06, all *p*s < .001). Additionally, German narrations led to a significant decrease in activation (raSTS: *t*(22) = -3.50, *p* = .002, laSTS: *t*(22) = -1.97, *p* = .06), whilst for all other conditions, PSC was at baseline. Thus, aSTS showed a similar effect of comprehensibility as SI-pSTS. Finally, these main effects were qualified by a significant Language × Condition interaction. For bilateral aSTS, PSC was greater for conversations compared to narrations for both English and German stimuli (all *t*s(22) > 3.03, *p* ≤ .006). Finally, right aSTS showed a significantly greater response to conversations compared to scrambled conversations (*t*(22) = 2.78, *p* = .01) for English stimuli only, suggesting sensitivity to the *coherence* of comprehensible interactions.

**Figure 4.**
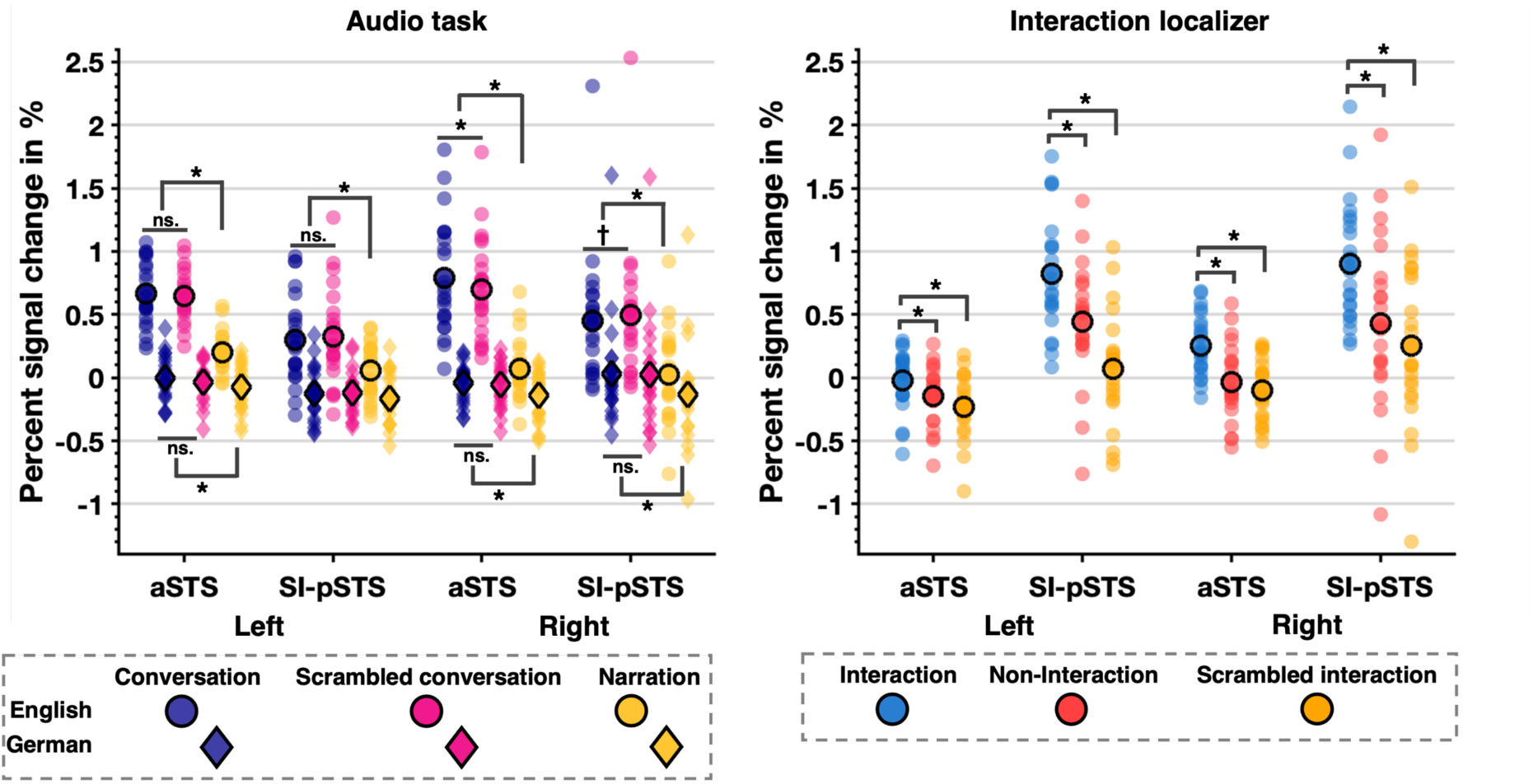
Percent signal change data displaying condition means (circle with bold edge) and data distribution for each aSTS and SI-pSTS ROI by hemisphere for audio task (left panel) and interaction localiser (right panel). Please note that pSTS data for the audio task is the same as in Figure 1 and for the interaction localiser is the same as Figure 2). Significant post hoc t-test results are marked by an asterisk (*: p < .0125 corrected, †: p < .05 uncorrected).

**Table 4.** ANOVA results of global aSTS peak ROI PSC analyses for main auditory task in Experiment 1

|  | Side | Main effect of Language |  |  | Main effect of Condition |  |  | Interaction effect |  |  |
| --- | --- | --- | --- | --- | --- | --- | --- | --- | --- | --- |
| | | *F* | *p* | $\eta_p^2$ | *F* | *p* | $\eta_p^2$ | *F* | *p* | $\eta_p^2$ |
| aSTS | L | 161.20 | <.001 | 0.88 | 55.84 | <.001 | 0.72 | 54.21 | <.001 | 0.71 |
|  | R | 74.56 | <.001 | 0.77 | 78.07 | <.001 | 0.78 | 75.66 | <.001 | 0.78 |

#### 3.1.6 What is the evidence for heteromodal social interaction processing in aSTS?

Further exploratory analyses were conducted to investigate whether a region sensitive to auditory social interactions identified using our main auditory experimental task would also be responsive to visual social interactions. As such, this analysis was a reversal of our main experimental hypothesis, i.e., how does the *auditorily* defined aSTS region respond to *visual* interactions, to explore whether social interactions are processed cross-modally within the social brain. Thus, bilateral aSTS was used to extract PSC from the interaction localiser.

Visual interactions only activated right aSTS significantly above baseline (*t*(22) = 5.03, *p* < .001), whereas non-interactions (*t*(22) = -0.61, *p* = .55) and scrambled interactions (*t*(22) = - 1.96, *p* = .06) were at or marginally below baseline. For left aSTS, all conditions were at (interactions, *t(22)* = -0.44, *p* = .66) or significantly below (non-interactions *t*(22) = -2.95, *p* < .01; scrambled interactions, *t*(22) = -4.56, *p* < .001) baseline. Paired sample t-tests comparing interactions with non-interactions as well as interactions with scrambled interactions found significantly greater responses to interactions bilaterally for aSTS (all *t*s(22) > 3.71, all *p*s ≤ .001); see Figure 4 right panel and Supplementary S7, Table S11).

Finally, comparison of interaction-selectivity in the auditory vs visual domain revealed the reverse pattern to SI-pSTS (see 3.1.3), greater auditory interaction selectivity (raSTS: M = 0.83, SE = 0.09, laSTS: M = 0.54, SE = 0.07) compared to visual interaction selectivity (raSTS: M = 0.29, SE = 0.05, laSTS: M = 0.12, SE = 0.07) (raSTS: *t*(22) = -7.07, *p* < .001, *d_rm_* = 0.36, laSTS: *t*(22) = -5.94, *p* < .001, *d_rm_* = 0.29).

#### 3.1.7 Summary

The main aim of this experiment was to investigate whether the interaction-sensitive SI-pSTS region is not only responsive to visual but also responsive to auditory interactions. Both univariate and multivariate ROI analyses suggest that bilateral SI-pSTS displays interaction-sensitivity to a broad contrast of *two speakers vs one speaker.* Univariate results also lend tentative support that right SI-pSTS exhibits interaction sensitivity *beyond* the number of speakers. This notion was corroborated more strongly using decoding analyses. Specifically, right SI-pSTS was the only region which could decode conversations vs. scrambled conversations, indicating that it also represents information about the meaningfulness of an auditory interaction. Unexpectedly, there were strong effects of language; bilateral TVA responded above baseline across both languages, whereas SI-pSTS was not driven by German stimuli. In contrast to our predictions, it seems likely that language comprehension was an important factor in some of our results. However, right SI-pSTS could discriminate between German conversations and narrations, suggesting that comprehension is not a pre-requisite when processing interactions at the level of speaker number. Finally, bilateral TVA also exhibited sensitivity to interaction based on the number of speakers, and unexpectedly, like SI-pSTS, left TVA also exhibited weak interaction sensitivity *beyond* the number of speakers, but only in univariate analyses.

Furthermore, this experiment explored (1) whether there was another brain region particularly sensitive to auditory interactions, and (2) whether visual and/or auditory interaction-sensitive regions may exhibit a heteromodal response profile. We used whole-brain group response to “find” a region in bilateral aSTS and explored its auditory and visual interaction sensitivity. Interestingly, right aSTS displayed a response profile characterized by greater sensitivity to auditory than visual interactions, whereas the right SI-pSTS showed the opposite pattern of greater sensitivity to visual compared to auditory interactions. Importantly, both regions showed sensitivity to interactive content *across* modality.

### 3.2 Experiment 2

This experiment was conducted as a small-scale follow-up study to address the unexpectedly strong language effects observed in Experiment 1. Here, participants were fluent in English but not German; thus, language comprehension might have driven PSC responses. For instance, SI-pSTS bilaterally was either at or below baseline for German conditions. This might be the result of listening to recordings in a language one does not comprehend in the context of a language you understand very well within the same run. A similar native vs unknown language comprehension effect has been found in prior work (Cotosck et al., 2021) using a target word detection task whilst listening to stories. On the other hand, language-specific acoustic differences (Mennen et al., 2012, see also Supplementary S4.2) might have driven some of these differences, particularly in the TVA. Experiment 2 set out to address this question with particular focus on the SI-pSTS by using the same stimulus set but testing German-English bilingual participants.

Please refer to Table 2 for ANOVA statistics, Supplementary S7, Table S7 for condition means, and Figure 5 for an illustration of the PSC results of Experiment 2.

**Figure 5.**
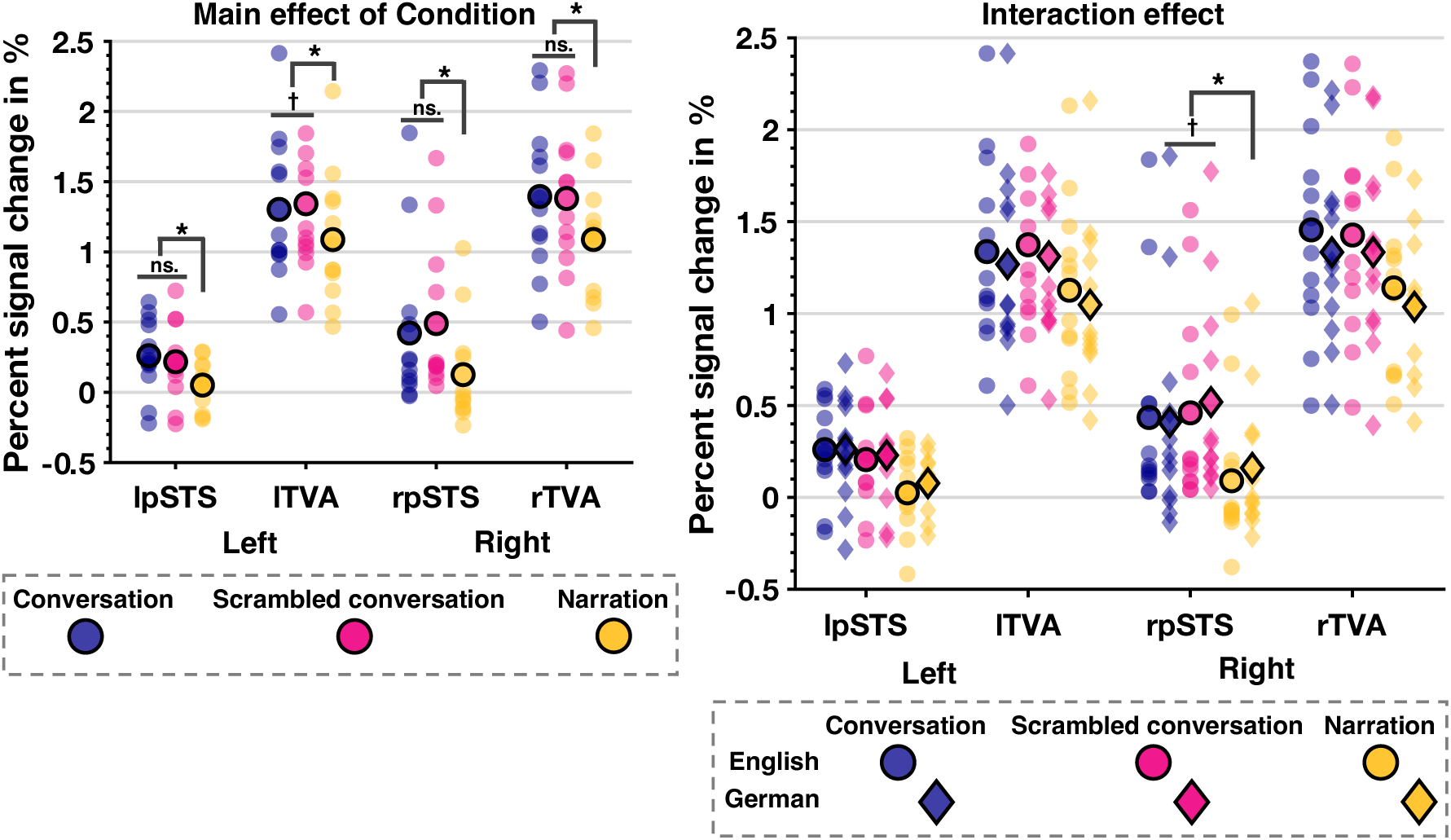
Experiment 2 PSC data illustrating the main effect of Condition (*: p < .017 corrected) and Language × Condition interaction effect (*: p < .0125 corrected, †: p < .05 uncorrected)

#### 3.2.1 Does SI-pSTS respond differently across languages when both are understood?

When both languages were comprehensible to participants, there was no main effect of language. Responses in bilateral SI-pSTS were similar for English and German stimuli. Indeed, PSC in bilateral SI-pSTS was significantly above baseline for both English and German conversations and scrambled conversations (all *t*(11) ≥ 2.36, all *p*s < .04). Thus, regardless of language, only narrations *did not activate* the SI-pSTS. Replicating Experiment 1, there *was* a main effect of Condition, driven by a difference between hearing two speakers vs. hearing only one. PSC was smaller for narrations compared to both conversations (rSI-pSTS: *t*(11) = 4.41, *p* < .001, *d_rm_* = 1.04, lSI-pSTS: *t*(11) = 3.66, *p* < .001, *d_rm_* = 0.25) and scrambled conversations (rSI-pSTS: *t*(11) = 6.35, *p* < .001, *d_rm_* = 0.80, lSI-pSTS: *t*(11) = 3.17, *p* < .001, *d_rm_* = 0.27) but not different between conversations and scrambled conversations. These main effects were qualified by a significant Language × Condition interaction in the right SI-pSTS only. PSC was greater for conversations compared to narrations for both English (*t*(11) = 5.23, *p* < .001, *d_rm_* = 0.41) and German stimuli (*t*(11) = 3.31, *p* = .007, *d_rm_* = 0.26). In contrast, response to *German* (*t*(11) = -2.38, *p* = .04, *d_rm_* = 0.18) but not English (*t*(11) = -0.59, *p* = .57) scrambled conversations was significantly greater than conversations, albeit at an uncorrected level only. This suggests some sensitivity to the *coherence* of interactions in the participants’ *native* language.

#### 3.2.2 How does the TVA response compare in this case?

As in Experiment 1, all conditions strongly activated bilateral TVA above baseline (all *t*(11) ≥ 7.69, all *p*s < .001). Bilaterally, although marginally in the left hemisphere, a significant effect of language remained even when participants understood both languages. Similarly, PSC was greater for English compared to German conditions. Further replicating Experiment 1, there was also a main effect of Condition. PSC was smaller for narrations compared to both conversations (rTVA: *t*(11) = 4.70, *p* < .001, *d_rm_* = 0.57, lTVA: *t*(11) = 4.82, *p* < .001, *d_rm_* = 0.52) and scrambled conversations (rTVA: *t*(11) = 4.53, *p* < .001, *d_rm_* = -0.18, lTVA: *t*(11) = 5.89, *p* < .001, *d_rm_* = 0.56). Finally, sustaining the unexpected finding from Experiment 1 of lTVA sensitivity to not only number of speakers but also coherence of conversation, responses to scrambled conversations were slightly but significantly greater than for intact conversations in left TVA only, albeit at an uncorrected level (*t*(11) = -2.61, *p* < .03, *d_rm_* = 0.56). No significant interaction effect emerged.

#### 3.2.3 Summary

Experiment 2 set out to test whether language comprehension may have driven some of the effects seen in Experiment 1. Testing bilingual participants revealed that when participants comprehended both languages, language effects disappeared in bilateral SI-pSTS whilst condition effects remained. In line with the results from Experiment 1, SI-pSTS responded more strongly to conversations compared to narrations. Crucially, right SI-pSTS was sensitive to the difference between German conversations and German scrambled conversations. This replicates Experiment 1 which found this difference for English stimuli. Taken together, these findings suggest right SI-pSTS is sensitive to meaningful auditory interactions, at least in participants’ native language. For bilateral TVA, language effects remained relatively stable with the participants’ non-native language resulting in greater activation. This might point to more effortful processing of the participants’ second language (Hasegawa et al., 2002). Like Experiment 1, left TVA showed a greater response to scrambled compared to intact conversations. Thus, across both experiments, left TVA was less responsive to meaningful conversations.

## 4. Discussion

Whilst everyday social interactions provide a rich multi-sensory experience, neuroimaging studies of social interaction perception have predominantly focused on the abundance of visual cues they provide. Conversely, not much is known about auditory interaction perception in the social brain. Combining univariate and multivariate analyses, we confirmed our key prediction that visual SI-pSTS exhibits heteromodal processing of social interactions. In contrast, although voice-selective TVA shows an unexpectedly similar response profile to auditory interactions, it is clearly a unimodal region. Specifically, both bilateral SI-pSTS and TVA were sensitive to interactive information in a broad contrast between two-speaker conversations and one-speaker narrations, in line with similar recent work focused on language processing (Olson et al., 2023). More importantly, right SI-pSTS and left TVA showed some weak sensitivity to auditory interactions when coherence of comprehensible (native language) conversations was manipulated. However, multivariate decoding analyses only corroborated this finding for right SI-pSTS, suggesting that the information represented in SI-pSTS voxel patterns is qualitatively different from that represented in TVA.

These findings are in line with previous results that put the broader pSTS region at the heart of heteromodal or even multimodal integrative processing of social information (Kreifelts et al., 2009; Lahnakoski et al., 2012; Robins et al., 2009; Watson et al., 2014; Wright et al., 2003). Indeed, regions along the pSTS show tuning to a variety of both visual and auditory social stimuli (Deen et al, 2015) and the pSTS is widely referred to as the ‘hub’ of the social brain because of its involvement across varied social tasks (e,g., Yang et al., 2015). However, much of the prior literature has investigated heteromodal processing in the context of social signals from individuals, making this study’s focus on the perception of social interactions relatively unique. Importantly, our data make clear that auditory interaction sensitivity in right SI-pSTS reflects more than tuning to voice stimuli in general. Indeed, the SI-pSTS region shows negligible sensitivity to vocal sounds in response to the voice localiser (see Figure 2). At the same time, response in SI-pSTS to visual and auditory interactive stimuli were not fully equivalent. Right SI-pSTS interaction-selectivity for visual stimuli was about 50% greater than for auditory stimuli. It could be that the nature of the interaction-region localiser might in part account for this. Essentially, we tested how SI-pSTS voxels sensitive to interaction information conveyed by human body- and biological motion cues responded to interaction cues conveyed by voice. Body and voice cues, however, are less strongly associated with each other compared to face and voice cues. Had we used stimuli that relied on facial cues of interaction in our localiser, we might have found a greater degree of correspondence between visual and auditory SI-pSTS response profiles. Indeed, heteromodal responses in the broader STS region to voices have previously been shown in conjunction with face stimuli (Deen et al., 2015; Deen et al., 2020; Watson et al., 2014), though not in the context of social interactions. As such, our approach is a strong test of whether SI-pSTS shows sensitivity to interactive information across modalities.

Nonetheless, the response profile of SI-pSTS to auditory interactions was more nuanced and less definitive than originally predicted. As a broad test of auditory interaction sensitivity, we expected that the mere presence of two speakers taking conversational turns would drive SI-pSTS activation regardless of language comprehension. However, testing monolinguals (Experiment 1) and bilinguals (Experiment 2) revealed that comprehension mattered. The SI-pSTS was only driven by the two-speaker conditions in monolingual participants’ native language, whereas this language effect was abolished in bilingual speakers. Nevertheless, in monolingual English speakers, MVPA analysis of voxel-patterns of SI-pSTS revealed that the two-speaker conditions could be discriminated from narrations even in the German condition. While this is perhaps not surprising, it does suggest that language comprehension is not a pre-requisite for representation of information; i.e., number of speakers, that clearly distinguishes auditory interactions from non-interactions within the SI-pSTS. When this distinction is less obvious however, interactive cues might well be derived through language comprehension. Indeed, monolingual participants listening to intact and scrambled conversations presented in their native language would be able to differentiate them based on detecting conversational coherence and presence of overall gist, whereas without comprehension, they would have to rely on subtle prosodic cues. We found that *right* SI-pSTS only distinguished between the two-speaker conditions when participants could access their meaning. Thus, language comprehension clearly mattered when extracting cues to conversational coherence. Notably, SI-pSTS lies in proximity to a bilateral brain network (including STG, STS, MTG and left IFG, see Yang 2019, Walenski 2019, Mar 2012; Bookheimer, 2002; Vigneau et al., 2011) implicated in higher-level discourse comprehension processes such as evaluation of global coherence, pragmatic interpretations, and text integration at the gist-level. However, right SI-pSTS’s overlap with this network is unclear. Although domain-general language processes designed to detect coherence could contribute to the response difference between intact and scrambled interactions, they cannot explain the drop in response to coherent narrations. Instead, right SI-pSTS might receive and integrate coherence or gist information from nearby language regions as a cue to evaluate interactiveness. Furthermore, although we found no support that SI-pSTS was sensitive to conversational flow in an unknown language, strong between-language effects may have overshadowed the potential to detect more subtle effects of prosody. Future studies investigating the role of SI-pSTS independent of language comprehension are needed to firmly establish its role when cues to interaction are harder to extract; e.g. using low-pass filtered muffled stimuli containing only prosodic but no lexico-semantic cues to interaction.

Unexpectedly, voice-selective TVA, especially in the left hemisphere, exhibited a similar response profile to right SI-pSTS. Firstly, a greater response to and decoding of conversations vs. narrations across languages, and secondly, a slightly greater response to scrambled conversations compared to conversations. Importantly, however, classification analyses in left TVA could not discriminate scrambled from intact conversations. Thus, it is unclear how distinctly (left) TVA responses could be attributed to pure interaction sensitivity. Indeed, it is possible that response difference between the one-speaker narration condition and the two speaker conditions (conversations, scrambled) in both TVA and SI-pSTS could partially be driven by these regions adapting to vocal quality or speaker identity in the narration condition. However, pSTS responsivity to voices is thought to reflect higher-level social process because individuals with lesions in pSTS are still able to discriminate between and recognise individual voices (Jiahui et al, 2017). In addition, we think it unlikely that adaptation can fully explain our effects in the SI-pSTS because we do not see strong differences between these conditions when participants don’t understand what is being said, making simple adaptation effects unlikely. However, as TVA is known to be involved in the spectro-temporal analysis of human vocal sounds (and speech) (Agus et al., 2017; Belin et al., 2002; Belin et al., 2000), adaptation to vocal quality or identity may partially explain our effects in this region. Similarly, previous research has found a significant positive association between mean F0 of speech and TVA activation (Wiethoff et al., 2008). More generally TVA is part of the STS/STG engaged in phonological language processing (Vigneau et al., 2006; Vigneau et al., 2011). Notably, due to the complexity and diversity of the stimuli used in this study, acoustic features of the stimulus set could not be as tightly controlled as we might have liked, which may drive some between-condition differences in TVA activation. Specifically, F0 was greater for conversations compared to narrations, and greater for English compared to German stimuli (see Supplementary S4.2). Thus, greater TVA activation to conversations compared to narrations, and English compared to German observed across both experiments could be at least partially explained by their corresponding differences in F0. However, as intact and scrambled conversations were matched on F0, these differences cannot explain higher left TVA response to scrambled conversations. Importantly, although scrambled and intact conversations contained identical sentences, they were not exact phonological equivalents as our scrambling process allowed a change in speaker. Thus, differential responses in left TVA might reflect sensitivity to speaker-dependent variations in phonation between conditions. Importantly, the above explanations would not apply to responses in right SI-pSTS, which is not known to be involved in phonological processes (Vigneau et al., 2006; Vigneau et al., 2011) or modulated by F0 (Wiethoff et al., 2008). In addition, between-language differences were no longer present in the SI-pSTS when participants understood both languages (Experiment 2). Altogether, our findings might motivate future work into auditory interaction perception in the brain to corroborate evidence on the unique role of right SI-pSTS using more tightly controlled stimuli that could additionally clarify the role of TVA.

Finally, we confirmed our prediction that TPJ, a social cognition region selectively engaged by mentalising processes (Saxe et al., 2009; Van Overwalle & Baetens, 2009), is neither driven by our stimuli nor sensitive to differences between conditions (see Supplementary S5 for details). TPJ can be engaged by auditory stimuli when participants are engaged in a mentalising task (Kandylaki et al., 2015; Saxe et al., 2009). However, while some prior work has suggested that TPJ is involved in processing social interactions (e.g., Canessa et al., 2012; Centelles et al.,2011), it is likely that the region is involved only when mentalising is required (Masson & Isik, 2021; Walbrin et al., 2018) which was not the case in our task. Indeed, we took care to select stimuli that did not imply nor require mentalising. Similarly, our whole-brain analysis suggests that occipital and temporal areas outside the STS, including EBA, are not involved in auditory interaction perception. This makes sense, as ‘early’ social perception regions like EBA are not usually considered to be heteromodal or, indeed, particularly responsive to auditory stimuli (Beer et al., 2013). Instead, our exploratory whole-brain analysis identified an area in right anterior STS, close to the temporal poles, which showed sensitivity to interactive information both through greater activation to two speakers than to one and higher response to comprehensible conversations compared to scrambled conversations (Experiment 1). While this region’s responsiveness to auditory interactions (and perhaps to interactive information in general) needs to be replicated (though see Olsen et al, 2023), this is a particularly intriguing finding in light of prior work that highlight the dorsolateral anterior temporal lobe (ATL) as a region that may be involved in social semantics (Arioli et al, 2021; Lin et al., 2018; Zhang et al., 2021; but see also Balgova et al., 2022; Binney et al., 2016; Binney & Ramsey, 2020 for a broader perspective regarding the role of the ATL in social cognition). Furthermore, in the context of speech comprehension, it has been suggested that the meaning of speech is processed in bilateral anterior temporal cortex, including aSTS (e.g., Mitchell et al., 2003; Scott et al., 2000; Scott et al., 2009, for a review see Price, 2012 . Thus, right aSTS might be particularly involved in the semantic analysis of auditory interactions, given that its activation was not driven by comprehensible narrative stimuli.

## Conclusion

Our results present initial evidence that SI-pSTS, initially defined visually, is also sensitive to interactive cues presented in the auditory domain. In other words, this region is characterised by a heteromodal response profile that appears to be particularly sensitive to social interactions. Future research is needed to both replicate these novel findings and to look beyond the number of speakers and interaction coherence to investigate whether SI-pSTS codes for other auditory cues involved in understanding social interactions, for instance subtle prosodic cues or interactional turn-duration. In addition, our results may motivate future work to determine how SI-pSTS integrates multimodal audio-visual social interaction information to inform our understanding of highly naturalistic everyday life social interactions. Finally, this initial work also prompts further research into the role of aSTS regions in social interaction perception more broadly and in conversation/language-based interactions specifically.

## Supplementary materials

### S1. Behavioural results: Catch trial detection during the main auditory task

In Experiment 1, on average, participants correctly detected 10.08 catch trials per included run (∼84%). Participants correctly detected significantly more (*t*(*22*) =7.72, *p* < .001) catch trials per run for recordings in English (M=5.29, SD = 0.29) compared to German (M=4.80, SD = 0.33).

Due to technical issues with the response device, behavioural data is missing completely for four participants, and for one out of seven runs for another participant (remaining data included) in Experiment 2. On average, participants correctly detected 10.12 catch trials per included run (∼84.5%). Participants correctly detected significantly more (*t*(*7*) =3.20, *p* = .02) catch trials per run for recordings in English (M=5.22, SD = 0.29) compared to German (M=4.91, SD = 0.27).

### S2. Details of audio recordings and audio processing

All audio was recorded directly into Logic Pro X (Release 10.4.2, Apple Inc.) audio processing software using one Shure SM58 and one Shure SM57 microphone plugged into a Zoom H6n Audio Interface. During conversations, actors’ voices were recorded simultaneously using the two microphones.

The separate audio channels were normalised and processed using the Logic Pro X Channel Equaliser. To remove unwanted background noise, the audio was first band-pass filtered (filtering signal below 61Hz and above 12000Hz) using the ‘Spoken Word Vocal EQ’ pre-set, and then filtered using two instances of RX iZotope RX 7 Voice De-Noise. Next, the audio was compressed using the Logic Pro X Compressor. To remove any colour added by the noise filters or the compressor, the audio was equalised again using the ‘Spoken Word Vocal EQ’. Finally, the audio was sliced into the individual vocal samples using the Logic Pro X editing suit and rendered at a sample rate of 44100Hz and at 16-bit resolution. The final set of stimuli were root mean square amplitude-normalized and a custom Sensimetrics earphones EQ filter was applied.

### S3. Stimulus selection process

A total of 108 scripted conversations were recorded with a male and a female speaker pair for each language. Each conversation was recorded twice so that both speakers took the role of speaker A and speaker B. Two samples were taken for each recording to select the best quality one afterwards, resulting in 432 recordings per language & speaker pair. These recordings were first assessed for initial quality by two native English speakers regarding how natural they sounded. As a result of this “first-pass” selection, 69 English conversations (=138 recordings for speaker A & B total) were selected to be rated for naturalness, valence, interactiveness, and mental imagery on a 5-pt Likert scale (4 raters each rated half the set of conversations). Based on the highest ratings for naturalness and interactiveness, the final set of 26 English conversations was selected, scrambled conversations were generated, and then the final stimuli were rated again (see S4.3). German conversations were reduced to a set of 73 after a “first-pass” for initial quality by one native German speaker and one speaker with a good working knowledge of German. The final set of 26 German conversations was chosen to be *not* translated equivalents of the English conversations to minimise the occurrence of cognate words.

A total of 36 scripted narrations were recorded with two male and two female speakers for each language, resulting in 288 recordings total. The final set of 26 English narrations was based on ratings for naturalness (see S4.3) after excluding scripts that were deemed to potentially evoke theory-of-mind processes. Due to the smaller number of recorded narrations, German narrations included 16 stimuli that were translated English narrations.

Examples as well as the scripts of the final set of stimuli can be found as an Excel file on https://osf.io/4xedj/.

### S4. Final stimulus set characteristics

#### S4.1 Duration, word, and letter count

Per condition, English and German stimulus sets were matched for mean stimulus duration (Conversations: *t*(27) = 0.57, *p* = .57, scrambled conversations: *t*(27) = -0.07, *p* = .95, narrations *t*(27) = 0.76, *p* = .45). To achieve this, English and German stimulus sets were well matched on the level of letters per stimulus (*t*(25) = -0.66, *p* = .52), whilst they differed on the mean number of words per stimulus (*t*(25) = 6.02, *p* < .001), due to word length differences between the two languages. Thus, German stimuli contained fewer words with more letters to match English stimuli with more words but fewer letters.

**Table S1.** Total and mean stimulus duration, as well as mean letter and word count per stimulus by experimental condition for the final stimulus set

|  |  | **Total**<br>**duration in s** | **Mean stimulus**<br>**duration in s** | **Letter count**<br>**per stimulus** | **Word count per**<br>**stimulus** |
| --- | --- | --- | --- | --- | --- |
| Conversation | English | 211.55 | 7.56 (0.17) | 107.35 (2.09) | 26.65 (0.36) |
|  | German | 207.46 | 7.41 (0.12) | 107.77 (1.51) | 24.77 (0.36) |
| Scrambled<br>conversation | English | 211.77 | 7.56 (0.16) | 106.35 (1.75) | 26.42 (0.38) |
|  | German | 206.98 | 7.39 (0.13) | 107.12 (1.60) | 24.50 (0.33) |
| Narration | English | 247.05 | 8.82 (0.14) | 106.23 (1.62) | 26.08 (0.39) |
|  | German | 247.41 | 8.84 (0.15) | 108.04 (1.73) | 24.65 (0.32) |

#### S4.2 Mean pitch (F0)

A Language × Condition repeated-measures ANOVA revealed that mean F0 was greater for English compared to German stimuli (*F*(1,27) = 6.29, *p* = 0.02, *η_p_*^2^ = 0.19). Additionally, the mean F0 of the two-speaker conditions (conversations, scrambled conversations) was greater than the mean F0 of narration stimuli (main effect of condition: *F*(2,54) = 43.92, *p* < 0.001, *η_p_*^2^ = 0.62). There was no significant interaction effect, *F*(2,54) = 2.42, *p* = 0.10, *η_p_*^2^ = 0.08.

**Table S2.** Mean pitch (F0 in Hz) and standard deviation of stimulus set by condition and gender

| Language | Speaker gender | Conversation | Scrambled conversation | Narration |
| --- | --- | --- | --- | --- |
| English | Male | 159.51 (7.54) | 155.66 (10.07) | 134.18 (19.25) |
|  | Female | 217.14 (16.36) | 215.64 (16.69) | 191.81 (17.26) |
|  | *Across gender* | *188.33 (31.90)* | *185.65 (33.40)* | *163.00 (34.40)* |
| German | Male | 121.72 (7.77) | 118.76 (11.80) | 106.69 (12.26) |
|  | Female | 222.10 (7.29) | 226.24 (8.72) | 207.65 (7.22) |
|  | *Across gender* | *171.91 (51.64)* | *172.50 (55.60)* | *157.17 (52.35)* |

#### S4.3 Stimulus ratings

The final set of English narrations was rated by an independent sample of 8 raters (mean age = 30.8, SD = 5.08, 1 male) on perceived naturalness, valence, and the extent to which the narration evoked a mental image on a 5-point Likert scale (see Table S3). The final set of English conversations and scrambled conversations was rated by an independent sample of 27 raters (mean age = 22.3, SD = 5.1, 6 males) on perceived naturalness, valence, interactiveness (sounding like an interaction), closeness (relationship) of agents, as well as whether and to what extent the conversation evoked a mental image on a 5-point Likert scale.

**Table S3.** Mean and SD of English stimuli conditions

|  | Natural | Valence | % evoked<br>mental<br>image | Mental<br>image<br>strength | Interaction | Closeness<br>(relationship)<br>of agents |
| --- | --- | --- | --- | --- | --- | --- |
| **Conversations** | 3.94<br>(0.61) | 3.38<br>(0.35) | 74% | 3.59<br>(0.74) | 4.28<br>(0.53) | 3.24<br>(0.37) |
| **Scrambled**<br>**conversations** | 2.02<br>(0.80) | 2.79<br>(0.45) | 25% | 2.62<br>(0.77) | 1.94<br>(0.60) | 2.12<br>(0.66) |
| **Narrations** | 4.10<br>(0.42) | 3.47<br>(0.29) | 75% | 3.43<br>(0.66) | - | - |

Independent sample t-tests revealed that ratings did not differ between conversations and narrations; paired sample t-tests revealed that conversations were rated higher on all dimensions compared to scrambled conversations (see Table S4).

**Table S4.**
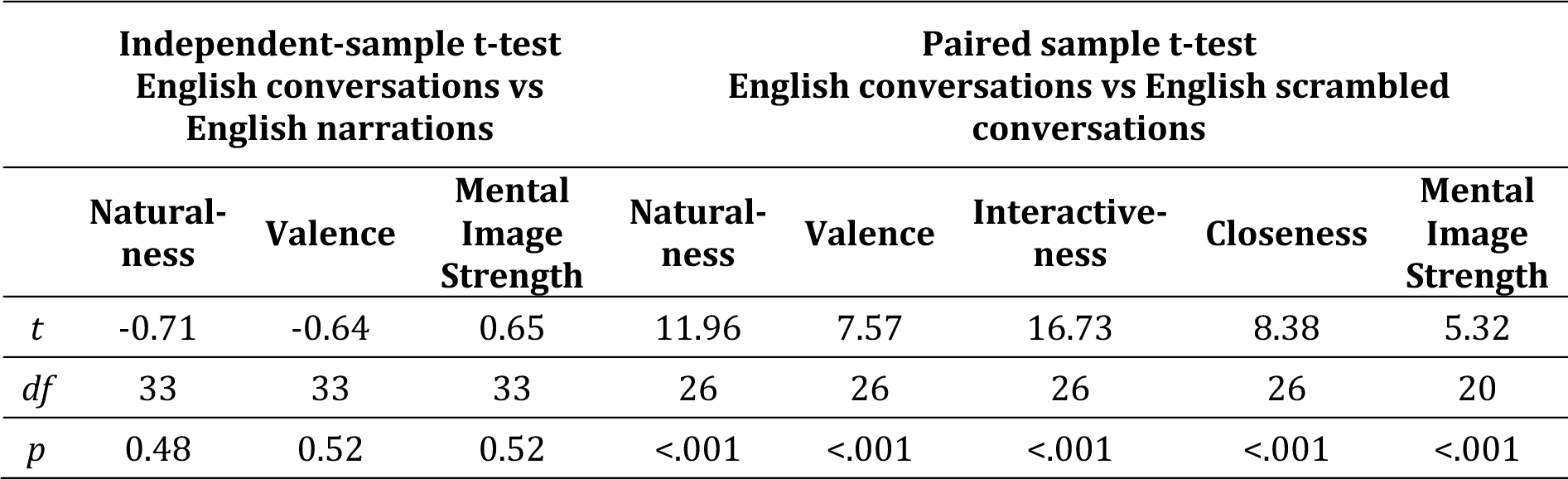
Rating data statistics

|  | Independent-sample t-test<br>English conversations vs<br>English narrations |  |  | Paired sample t-test<br>English conversations vs English scrambled<br>conversations |  |  |  |  |
| --- | --- | --- | --- | --- | --- | --- | --- | --- |
|  | Natural-<br>ness | Valence | Mental<br>Image<br>Strength | Natural-<br>ness | Valence | Interactive-<br>ness | Closeness | Mental<br>Image<br>Strength |
| *t* | -0.71 | -0.64 | 0.65 | 11.96 | 7.57 | 16.73 | 8.38 | 5.32 |
| *df* | 33 | 33 | 33 | 26 | 26 | 26 | 26 | 20 |
| *p* | 0.48 | 0.52 | 0.52 | <.001 | <.001 | <.001 | <.001 | <.001 |

### S5. TPJ as control ROI within the ‘social brain’

#### Method

Participants completed an additional localiser task (Jacoby et al., 2016) to identify the temporo-parietal junction (TPJ), which served as a control region within the ‘social brain’. TPJ was chosen based on previous findings that showed TPJ not to be sensitive to visual interactions across several stimulus types and paradigms (Isik et al., 2017; Masson & Isik, 2021; Walbrin et al., 2018; Walbrin & Koldewyn, 2019; Walbrin et al., 2020). In addition, the TPJ is very nearby the pSTS interaction region but not thought to be part of the voice-processing network. Thus, it was predicted that TPJ would not show sensitivity to the experimental conditions of the main auditory task. To localise the TPJ bilaterally, participants watched the short (5:49 minutes) Pixar animation short film ‘Partly Cloudy’ (2009) which has been found to reliably evoke responses in the mentalizing selective TPJ. The film scenes were coded by event type (mentalizing, pain, social, and control; Jacoby et al., 2016) and the contrast mentalizing vs. pain was used to localise TPJ. The same procedures as described in the main text were used to define subject-specific TPJ ROIs bilaterally.

#### Results

One-sample t-test against zero confirmed that the main auditory task did not drive activation in TPJ above baseline, suggesting these stimuli did not drive activity in the TPJ. Bilaterally, English conversations resulted in PSC that were not significantly different than baseline (all *t*s(22) > -1.17, all *p*s > .26), whereas marginally significant (German conversations, all *t*s(22) > -2.07, all *p*s > .05) or significant (all other conditions, all *t*s(22) < -2.11, all *p*s < .05 ) TPJ deactivation was observed otherwise. ANOVA revealed no significant main effects of Language (*F*(1,22) = 0.11, *p* = .74, *η_p_*^2^ < 0.01) or Condition (*F*(1,22) = 2.04, *p* = .14, *η_p_*^2^ = 0.09), nor a significant interaction (*F*(1.48,32.53) = 1.78, *p* = .18, *η_p_*^2^ = 0.08) in right TPJ. For left TPJ, there were no main effects of Language (*F*(1,22) = 0.15, *p* = .71, *η_p_*^2^ < 0.01) or Condition (*F*(1.40,30.77) = 2.56, *p* = .11, *η_p_*^2^ = 0.10), however, there was a significant interaction (*F*(1.38,30.31) = 5.46, *p* = .02, *η_p_*^2^ = 0.20). Explorative (uncorrected) post-hoc t-tests revealed that this was driven by a significant effect of language for narrations only: English narrations deactivated the region more than German narrations (*t*(22) = -2.47 , *p* = 0.02). Furthermore, English conversations deactivated the region significantly less than narrations (*t*(22) = 2.20 , *p* = 0.04).

**Table S5.**
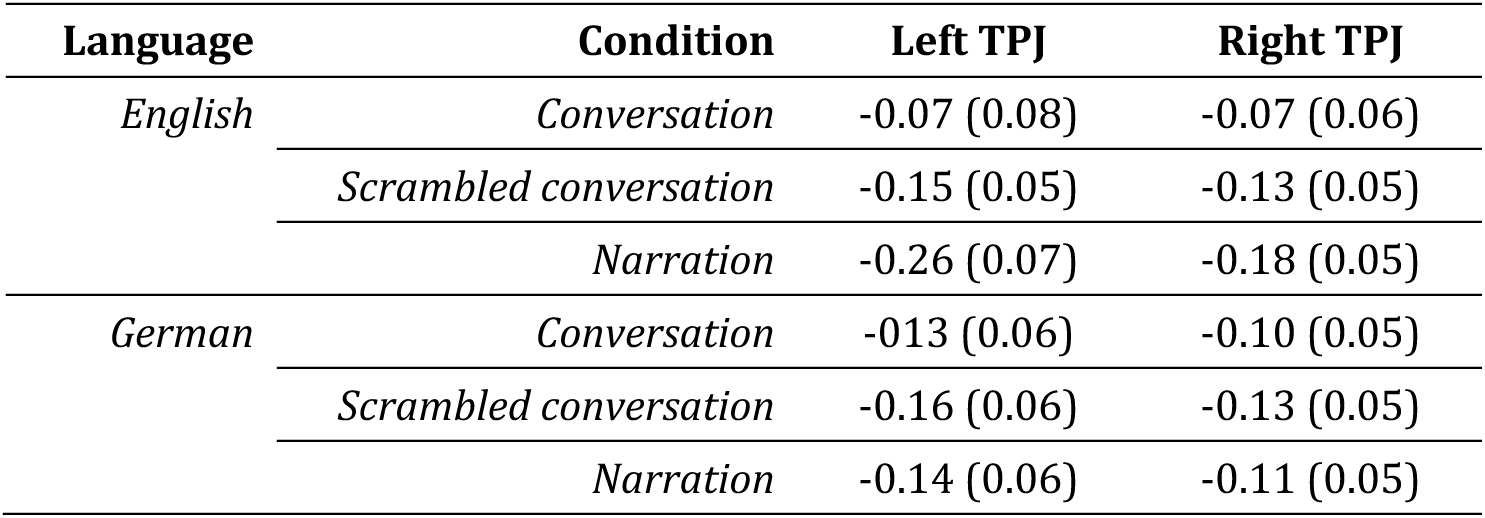
Bilateral TPJ PSC means (SE)

### S6. General ROI information

**Table S6.** MNI-space coordinates (based on group-level localiser task contrasts) used as centre for sphere creation identified at an uncorrected threshold of p < .01

| Region | Defining task | Defining contrast | Left | Right |
| --- | --- | --- | --- | --- |
| TPJ | Mentalizing localiser | Mentalizing > Pain | [-42 -68 40] | [48 -62 36] |
| SI-pSTS | Interaction localiser | Interaction > Non-Interaction | [-48 -60 12] | [52 -48 12] |
| TVA | Voice localiser | Vocal > Non-vocal sounds | [-60 -24 0] | [60 -24 0] |
| aSTS | Main audio task | Language × Condition interaction (F-contrast) | [-50 0 -24] | [52 2 -24] |

**Figure S1.**
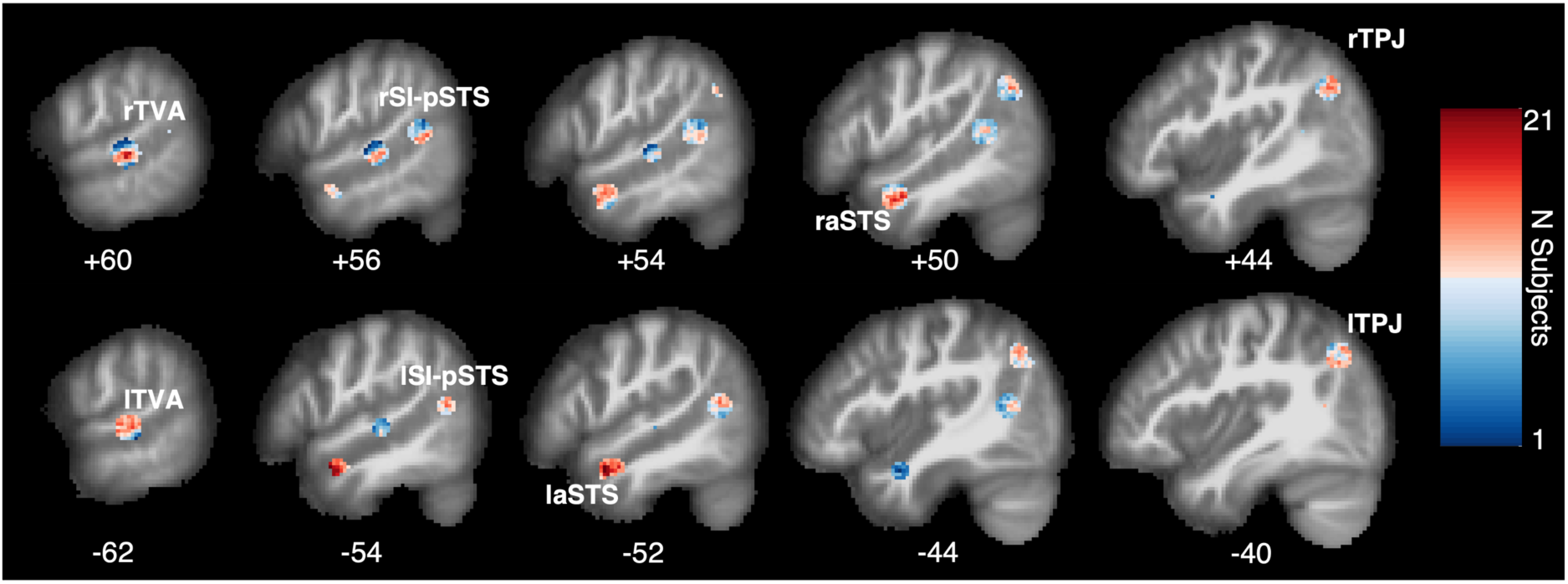
Sagittal view heatmap of subject-specific aSTS, TVA, SI-pSTS, and TPJ ROIs’ overlap for right (top row) and left (bottom row) hemisphere. x-coordinate in MNI space shown below each slice. Figure created using bspmview toolbox (DOI: 10.5281/zenodo.168074).

### S7. Additional data tables for Experiment 1 and Experiment 2

**Table S7.** SI-pSTS and TVA PSC means (SE) for all experimental conditions for both experiments

| Language | Condition | Experiment 1 |  |  |  | Experiment 2 |  |  |  |
| --- | --- | --- | --- | --- | --- | --- | --- | --- | --- |
|  |  | SI-pSTS |  | TVA |  | SI-pSTS |  | TVA |  |
|  |  | L | R | L | R | L | R | L | R |
| English | Conversation | 0.30<br>(0.07) | 0.45<br>(0.10) | 1.83<br>(0.13) | 2.01<br>(0.12) | 0.26<br>(0.07) | 0.43<br>(0.17) | 1.34<br>(0.15) | 1.45<br>(0.17) |
|  | Scrambled conversation | 0.33<br>(0.08) | 0.50<br>(0.11) | 1.88<br>(0.12) | 2.00<br>(0.12) | 0.21<br>(0.08) | 0.46<br>(0.16) | 1.38<br>(0.16) | 1.43<br>(0.16) |
|  | Narration | 0.06<br>(0.05) | 0.03<br>(0.08) | 1.50<br>(0.11) | 1.59<br>(0.12) | 0.02<br>(0.06) | 0.09<br>(0.11) | 1.13<br>(0.14) | 1.14<br>(0.13) |
| German | Conversation | -0.12<br>(0.04) | 0.03<br>(0.08) | 1.55<br>(0.11) | 1.70<br>(0.11) | 0.26<br>(0.09) | 0.41<br>(0.17) | 1.27<br>(0.15) | 1.33<br>(0.15) |
|  | Scrambled conversation | -0.12<br>(0.04) | 0.03<br>(0.09) | 1.52<br>(0.10) | 1.67<br>(0.11) | 0.23<br>(0.08) | 0.52<br>(0.16) | 1.31<br>(0.15) | 1.34<br>(0.15) |
|  | Narration | -0.16<br>(0.04) | -0.13<br>(0.08) | 1.31<br>(0.10) | 1.50<br>(0.12) | 0.08<br>(0.05) | 0.16<br>(0.11) | 1.05<br>(0.14) | 1.04<br>(0.11) |

**Table S8.** Bilateral PSC means (SE) for interaction localiser (Experiment 1 only)

| Condition | SI-pSTS |  | TVA |  | aSTS |  |
| --- | --- | --- | --- | --- | --- | --- |
|  | L | R | L | R | L | R |
| Interaction | 0.82<br>(0.10) | 0.90<br>(0.10) | -0.12<br>(0.06) | -0.21<br>(0.07) | -0.02<br>(0.05) | 0.26<br>(0.05) |
| Non-Interaction | 0.44<br>(0.10) | 0.43<br>(0.14) | -0.13<br>(0.05) | -0.21<br>(0.06) | -0.14<br>(0.05) | -0.04<br>(0.06) |
| Scrambled interaction | 0.07<br>(0.09) | 0.25<br>(0.13) | -0.16<br>(0.05) | -0.23<br>(0.07) | -0.23<br>(0.05) | -0.10<br>(0.05) |

**Table S9.** Bilateral PSC means (SE) for bilateral SI-pSTS responses to voice localiser (Experiment 1 only)

| SI-pSTS |  |  |
| --- | --- | --- |
| Condition | L | R |
| Vocal sounds | 0.01 (0.07) | 0.25 (0.12) |
| Non-vocal sounds | -0.06 (0.04) | 0.02 (0.06) |

**Table S10.**
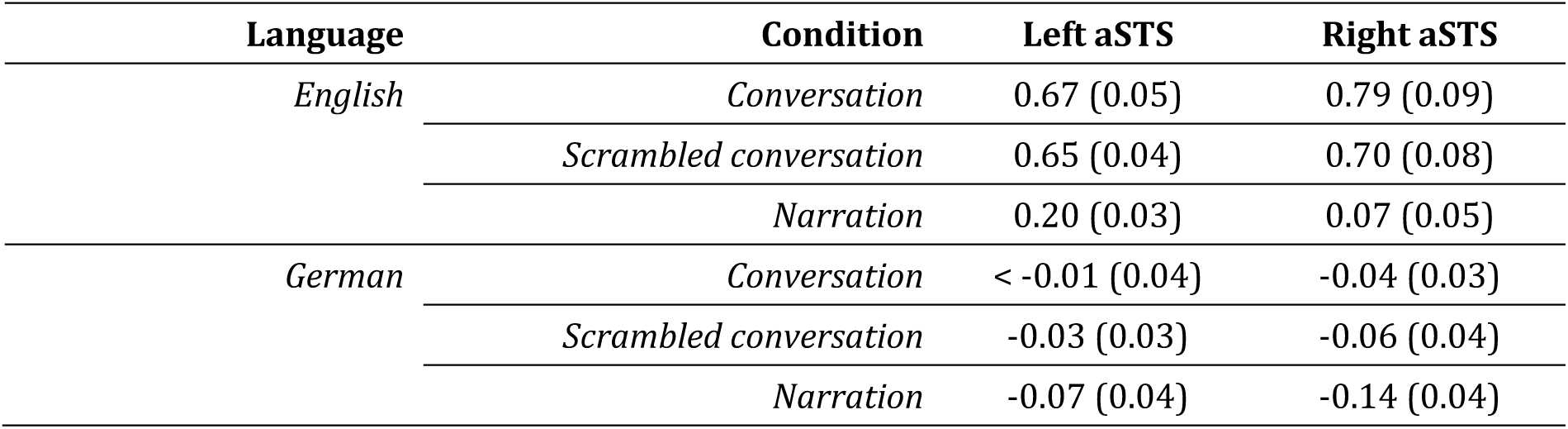
Bilateral aSTS PSC means (SE) for auditory task (Experiment 1 only)

| Language | Condition | Left aSTS | Right aSTS |
| --- | --- | --- | --- |
| English | Conversation | 0.67 (0.05) | 0.79 (0.09) |
|  | Scrambled conversation | 0.65 (0.04) | 0.70 (0.08) |
|  | Narration | 0.20 (0.03) | 0.07 (0.05) |
| German | Conversation | < -0.01 (0.04) | -0.04 (0.03) |
|  | Scrambled conversation | -0.03 (0.03) | -0.06 (0.04) |
|  | Narration | -0.07 (0.04) | -0.14 (0.04) |

**Table S11.**
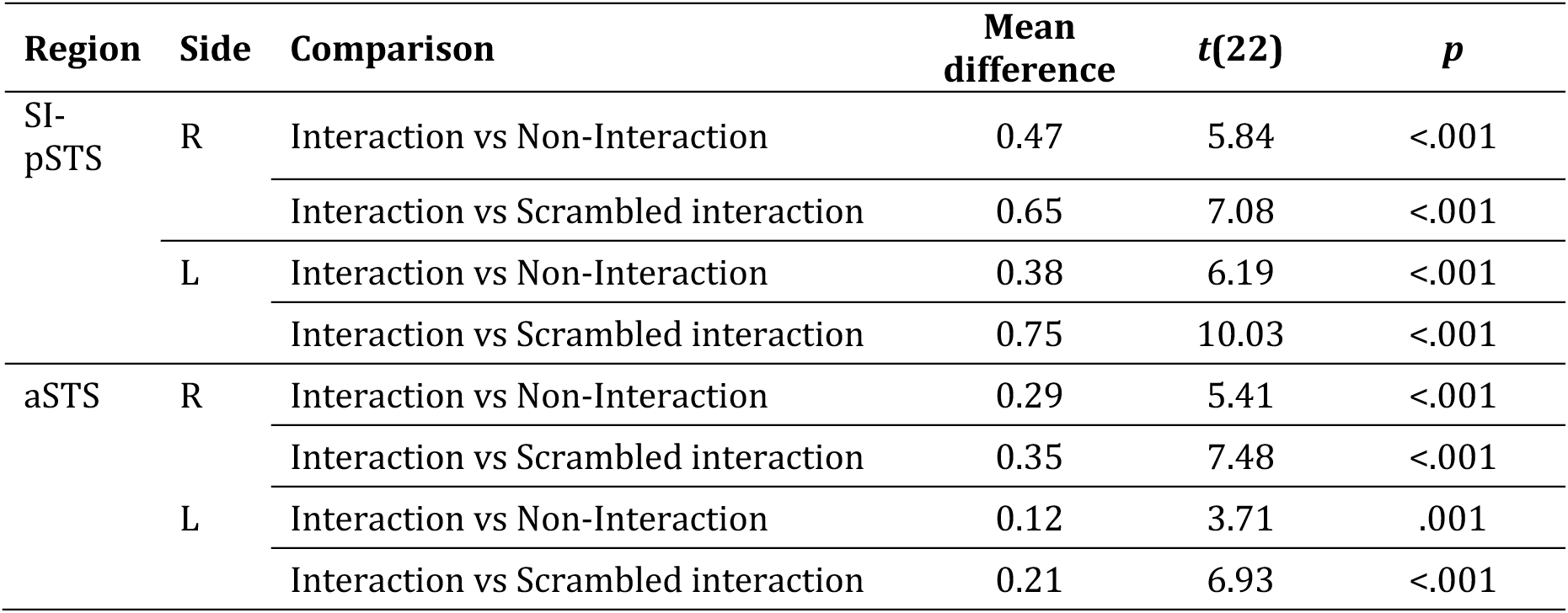
Results of paired sample t-tests for aSTS and SI-pSTS PSC comparison of interaction localiser conditions (Experiment 1 only)

| **Region** | **Side** | **Comparison** | **Mean difference** | **t(22)** | **p** |
| --- | --- | --- | --- | --- | --- |
| SI-pSTS | R | Interaction vs Non-Interaction | 0.47 | 5.84 | <.001 |
|  |  | Interaction vs Scrambled interaction | 0.65 | 7.08 | <.001 |
|  | L | Interaction vs Non-Interaction | 0.38 | 6.19 | <.001 |
|  |  | Interaction vs Scrambled interaction | 0.75 | 10.03 | <.001 |
| aSTS | R | Interaction vs Non-Interaction | 0.29 | 5.41 | <.001 |
|  |  | Interaction vs Scrambled interaction | 0.35 | 7.48 | <.001 |
|  | L | Interaction vs Non-Interaction | 0.12 | 3.71 | .001 |
|  |  | Interaction vs Scrambled interaction | 0.21 | 6.93 | <.001 |

## Author contributions

**Julia Landsiedel**: Conceptualization, Methodology, Investigation, Formal analysis, Project administration, Visualization, Writing – Original Draft, Writing – Review & Editing; **Kami Koldewyn**: Conceptualization, Funding acquisition, Methodology, Investigation, Supervision, Project administration, Writing – Original Draft, Writing – Review & Editing

## Acknowledgements

We are grateful to Corinne Voigt-Hill for her help with data collection as well to members of the Social Neuroscience and Cognition group and Bangor Imaging Group at Bangor University for general feedback, helpful discussion, and suggestions throughout the research process.

